# Being spontaneous has its costs! Characterization of spontaneous phage □D5-resistant mutants of *Dickeya solani* strain IPO 2222

**DOI:** 10.1101/2023.01.31.526416

**Authors:** Daryna Sokolova, Anna Smolarska, Przemysław Bartnik, Lukasz Rabalski, Maciej Kosinski, Magdalena Narajczyk, Dorota M. Krzyzanowska, Magdalena Rajewska, Inez Mruk, Paulina Czaplewska, Sylwia Jafra, Robert Czajkowski

## Abstract

Lytic bacteriophages able to infect and kill *Dickeya* spp. can be readily isolated from virtually all *Dickeya* spp.-containing environments, yet little is known about the selective pressure those viruses exert on their hosts. Here, we identified two spontaneous *D. solani* IPO 2222 mutants (0.8% of all obtained mutants), DsR34 and DsR207, resistant to infection caused by lytic phage vB_Dsol_D5 (ΦD5) that expressed a reduced ability to macerate potato tuber tissues compared to the wild-type, phage-susceptible *D. solani* IPO 2222 strain. Genome sequencing revealed that genes encoding: secretion protein HlyD (mutant DsR34) and elongation factor Tu (EF-Tu) (mutant DsR207) were altered in these strains. Both mutations impacted the proteomes of cells grown in both rich and minimal media, including the abundance of the cell envelope and transmembrane transport-associated proteins. Furthermore, features essential for the ecological success of these mutants in a plant environment, including their ability to use various carbon and nitrogen sources, produce plant cell wall degrading enzymes, ability to form biofilms, siderophore production, swimming and swarming motility and virulence *in planta* were assessed. Compared to the wild-type strain, *D. solani* strain IPO 2222, mutants DsR34 and DsR207 had a reduced ability to macerate chicory leaves and to colonize and cause symptoms in growing potato plants. The implications of the ΦD5 resistance on driving traits affecting the ecological performance of *D. solani* are discussed.

## Introduction

Lytic bacteriophages are the most abundant entities in the biosphere ^1^. They have been reported in virtually all environments inhabited by bacteria, where they are responsible for the global killing of 20 to 40% of host cells daily ^2, 3^. Due to their great killing potential, lytic viruses have been shown to exert strong selective pressure on their hosts ^4^. Consequently, they are well-known as one of the principal driving forces of bacterial adaptation and evolution ^5^. In addition, whereas bacteria are limited to one cell division per generation, a single bacterial cell infected by a single lytic phage virion can produce an average of more than a hundred progeny (daughter) viruses ^6^. Furthermore, these new phage particles can readily infect other bacterial cells in their vicinity, thus accelerating infection ^7^. As a result, lytic bacteriophages can rapidly overtake their bacterial hosts, leading to the rapid reduction of or even elimination of susceptible host cells ^8^. The potential for bacterial strains to persist in an environment replete with lytic viruses depends, therefore, on their ability to accumulate mutations that mediate their resistance to phage infection and, thus, the survival of such newly-resistant hosts under those conditions ^9^.

Although bacteria can interfere with viral infection at all stages of the phage-host interaction (including adsorption, DNA entry, replication, transcription and translation, capsid assembly, and release of progeny (daughter) phages) ^10^, phage-resistant mutants often avoid viral infections through mechanisms inhibiting virus adsorption to the host surface ^11^. This adsorption inhibition *via* spontaneous mutations in genes involved with the status of the cell surface has been best characterized at the genomic level in the model phage-host system of *Escherichia coli* and its phage T4 ^12, 13^. Likewise, several other studies have addressed this process in Gram-negative bacteria and their associated viruses ^14, 15^. Still, the knowledge about the relationship between the ecological costs of accumulating random mutations in the host genomes to achieve phage resistance, such as survival in adverse conditions, or the ability to remain virulent, is limited. Furthermore, the effect of phage-resistance mutations on other phenotypes of the resistant cells has not been well addressed. Finally, even though the occurrence of spontaneous phage resistance has been explored in several phage-bacterium systems, little is known about these processes taking place in plant pathogenic bacteria residing in their natural habitats, including phytopathogens present in agricultural settings. This subject is of particular interest given the diversity of environmental situations such bacteria face during their lives. Indeed, no such studies have addressed the spontaneous phage resistance and its ecological consequences in the important plant pathogenic bacteria, the Soft Rot *Pectobacteriaceae* (SRP: *Pectobacterium* spp. and *Dickeya* spp.) species ^16–18^.

Soft Rot *Pectobacteriaceae* are an excellent model for studying spontaneous phage resistance in the environment. *Pectobacterium* spp. and *Dickeya* spp. are considered among the ten most important agricultural plant pathogens and cause significant losses reaching up to 40% of crop production worldwide ^19^. Furthermore, *D. solani* is an emerging plant pathogen, causing disease symptoms in various crops and nonfood plants worldwide ^20^. This species was first reported in 2007 in potato ^21^, and has remained an important agricultural plant pathogen in most European countries ^22^ as well as in several agricultural regions outside Europe ^23–25^. The bacteria are prevalent in diverse ecological niches, including bulk soil and the rhizosphere of agricultural and natural soils, in surface and rainwater, and in host and non-host plants, as well as in insects ^22, 26, 27^. The large local population sizes that these bacteria achieve, especially in infected plants, facilitate epidemics of phage infection. Therefore, selective pressure associated with lytic bacteriophages in these settings is expected to lead to strong selection for phage-resistant mutants. The fitness of the mutants, thus, will depend not only on their virulence to plants but also on the particular environment in which bacteria are present and interact with the viruses. Likewise, SRP bacteria are often transferred between different ecological niches during their life cycles and are therefore exposed to new viral infections ^28^. However, the fitness costs the spontaneous phage-resistant SRP mutants pay in natural settings remain unclear.

This investigation sought to determine the extent to which spontaneous phage resistance and loss of virulence to plants are linked. Moreover, this study evaluated the extent to which spontaneous phage-resistant mutants of *D. solani* also exhibited other phenotypic alternations that could influence their ecological fitness compared to the wild-type, phage-susceptible strain. As a model, the study investigated the spontaneous phage resistance of mutants of strain IPO 2222 selected in the presence of lytic bacteriophage vB_Dsol_D5 (ΦD5). By demonstrating that spontaneous ΦD5-resistant *D. solani* IPO 2222 mutants exhibited decreased virulence, we addressed the hypothesis that resistance to viral infection negatively impacts bacterial phenotypes important during interactions with plants, thus lowering their fitness and decreasing competition in such environments.

## Results

### Selection of *D. solani* phage-resistant mutants expressing reduced virulence *in planta*

A total of 250 individual spontaneous ΦD5-resistant *D. solani* mutants were obtained in this study. All were confirmed to be *D. solani* IPO 2222 based on ERIC-PCR (data not shown). However, only 2 phage-resistant isolates (0.8% of all obtained and analyzed mutants, named DsR34 and DsR207) exhibited significantly reduced maceration of potato tubers compared to the WT strain (Fig. 1). These two phage-resistant mutants were chosen for further analysis. The selected ΦD5-resistant *D. solani* mutants expressed statistically significantly reduced ability to cause maceration symptoms on chicory leaves compared with the WT strain (Fig. 2). Likewise, in two separate experiments in which potato plants grown in potting soil were exposed to the strains with a soil drench, few plants inoculated with mutants DsR34 and DsR207 (0 to 20%) in both experiments exhibited any disease symptoms. In contrast, 80 to 90 % of the plants inoculated with the wild-type strain developed typical blackleg symptoms leading to the eventual death of the inoculated plants in both experiments. As expected, in both experiments, no symptoms were observed in control plants inoculated only with sterile Ringer’s buffer (Fig. 3A). In addition to the significant reduction in disease incidence in plants inoculated with the two phage-resistant mutants compared to the wild-type strain, the severity of infection, measured as the number of viable bacterial cells isolated from macerated plant tissue, was significantly reduced. Although the population size of the WT strain differed statistically between individual infected plants when measured 14 days after inoculation, the bacteria were recovered from the majority of inoculated plants at densities ranging from 10^3^ to 10^4^ CFU g^-1^ of the stem tissue (Fig. 3B). In contrast, DsR34 and DsR207 were seldom recovered, being found both in a much lower proportion of plants than the WT strain, and at a lower population size in the stem in those few plants in which it was recovered at all. Viable cells of DsR34 were detected in only 3 plants out of 20 inoculated (1 plant in experiment 1 and 2 plants in experiment 2). In contrast, cells of DsR207 were recovered in only 4 of 20 plants inoculated plants (2 plants in experiment 1 and 2 plants in experiment 2) expressing typical infection symptoms. The average population size of the phage-resistant mutants recovered from plants in both experiments were between 10^2^ to 10^3^ CFU g^-1^ of the stem tissue. *D. solani* was not detected in the stems of non-inoculated negative control plants (Fig. 3B).

**Figure 1.**
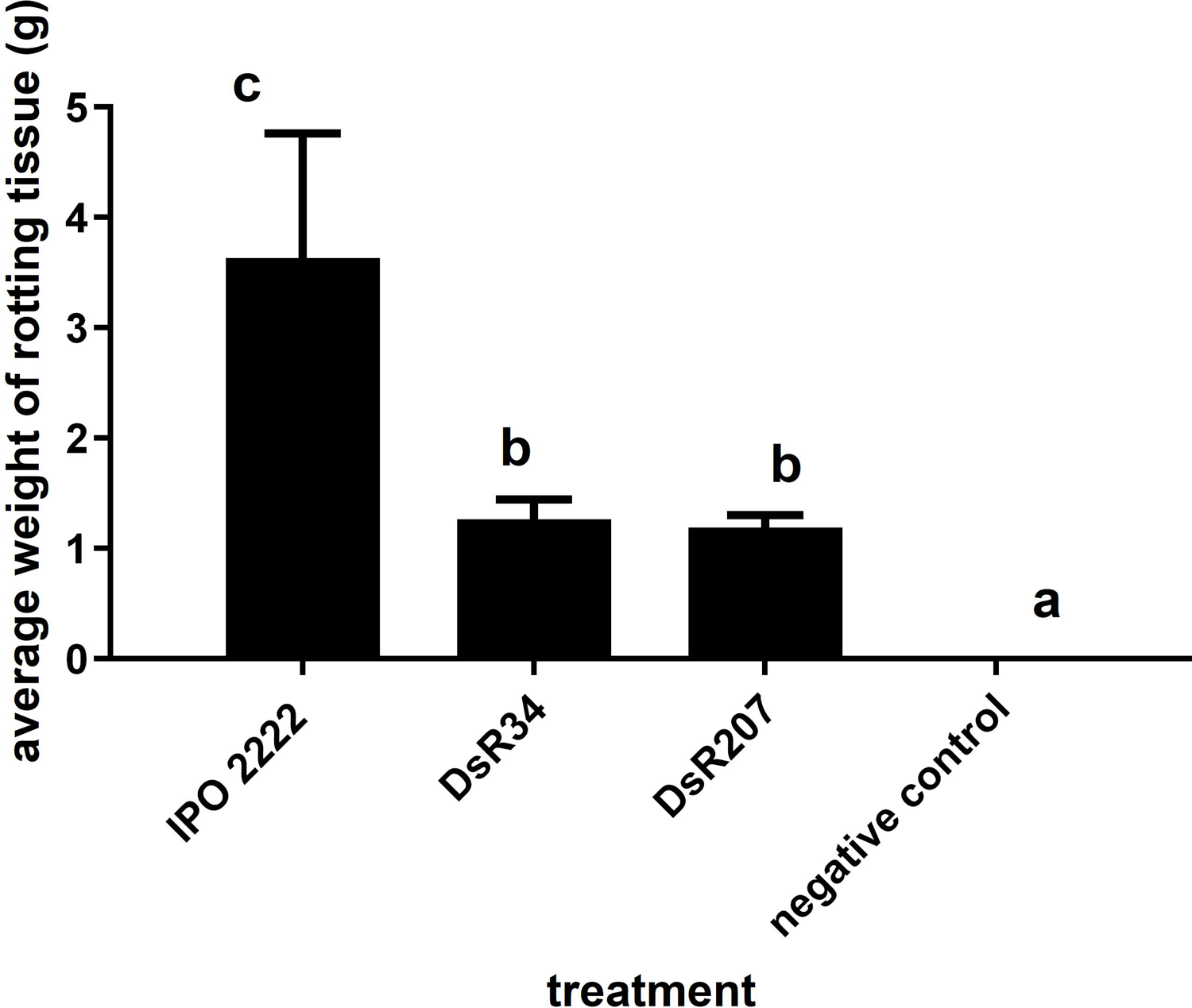
Ability of ΦD5-resistant *D. solani* mutants DsR34 and DsR207 to cause maceration (rotting) of potato tuber tissue. Quantitative determination of the average weight of the rotting tuber tissue (in grams) collected after 72 h incubation at 28 °C under humid conditions. Per mutant, five individual potato tubers were inoculated in two independent experiments (n=10). IPO 2222 wild-type strain was used as a positive control. Tubers inoculated with sterile demineralized water served as the negative control. Results were considered significant at p=0.05, and pairwise differences were acquired with the use of the *t*-test. The means not sharing the same letters above each bar vary. Error bars represent standard deviation (SD).

**Figure 2.**
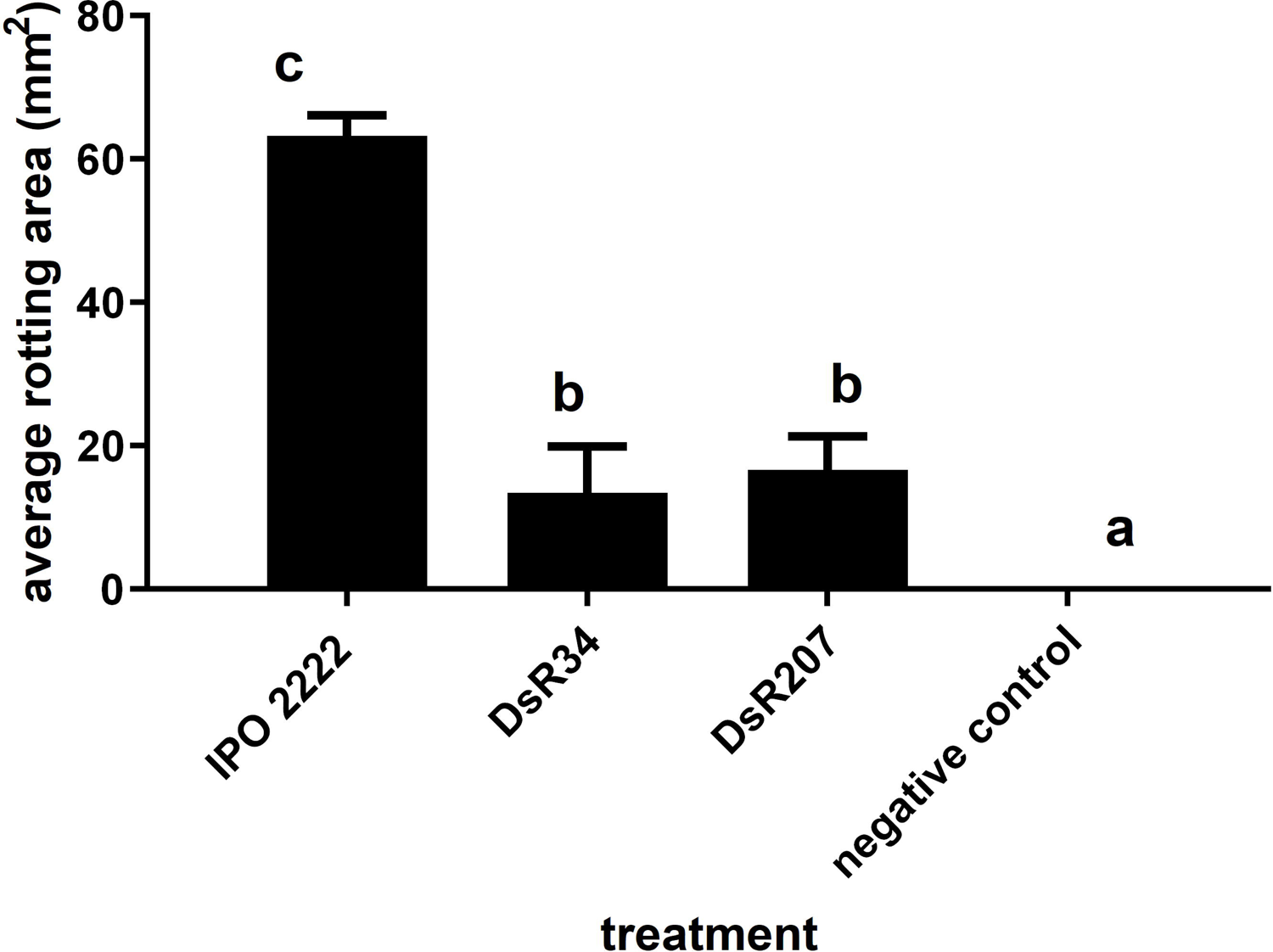
Ability of ΦD5-resistant *D. solani* mutants DsR34 and DsR207 to cause maceration (rotting) of chicory leaves. Quantitative determination of the average rotting area (in mm^2^) after incubation at 28 °C in a humid box. Per mutant, five individual chicory leaves were inoculated in two independent experiments (n=10). Results were considered significant at p=0.05, and pairwise differences were obtained using the *t*-test. The means not sharing the same letters above each bar vary. Error bars represent standard deviation (SD).

**Figure 3.**
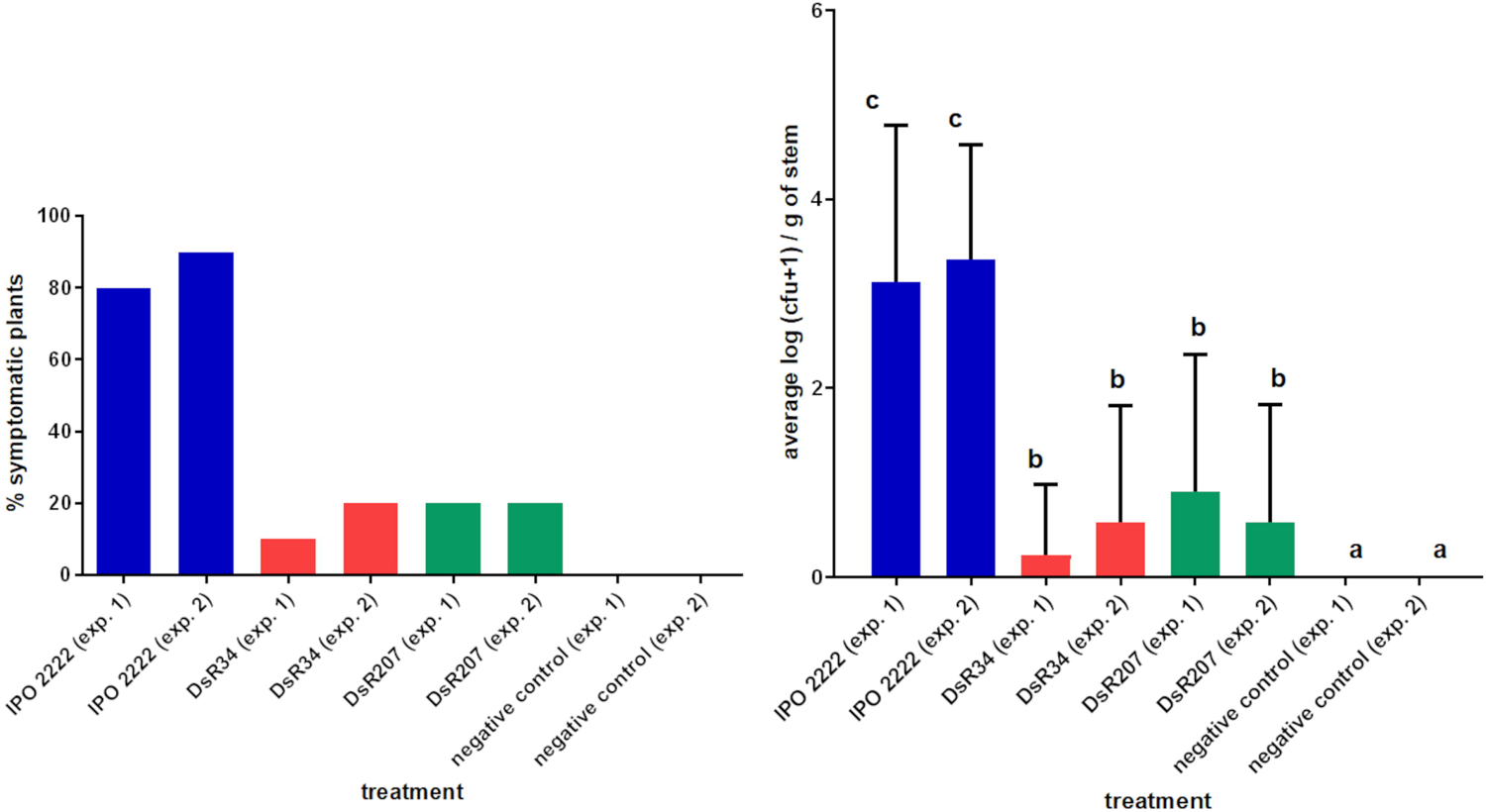
Ability of ΦD5-resistant *D. solani* mutants DsR34 and DsR207 to colonize and cause symptoms in potato plants cv. Kondor grown in potting compost under phytochamber conditions. Plants were grown for 14 days for rooting and shoot developments. Rooted and developed plants were inoculated with *D. solani* IPO 2222 wild-type strain or ΦD5-resistant DsR34 and DsR207 mutants (n=10 per treatment) by application of 50 ml of bacterial suspension (ca. 10^8^ CFU ml^-1^) in sterile Ringer’s buffer directly to the soil surrounding stem bases of each plant. As a negative control, the soil was treated with sterile Ringer’s buffer instead of the bacterial suspension. After two weeks, the samples were inspected for symptoms and bacterial populations inside stems. **A** – Percentage of symptomatic plants 14 days post inoculation (DPI) in both experiments. **B** – Population size of bacterial strains within stems of potato plants after introducing the pathogen into the soil in both experiments. Results were considered significant at *p* = 0.05, and the pairwise differences were obtained using the *t*-test. The means not sharing the same letters above each bar vary. Error bars represent standard deviation (SD).

### Sequencing of the genomes of *D. solani* phage-resistant mutants

The genomes of DsR34 and DsR207 mutants were sequenced to identify sites of putative genomic alternations mediating resistance to infection caused by ΦD5. Illumina sequencing generated 19 951 712 paired reads for *D. solani* mutant DsR34 and 194 888 056 paired reads for *D. solani* mutant DsR207. Obtained data was sufficient to achieve respective mean genome coverage of 602 for DsR34 and 586 for DsR207. Long-reads (Oxford Nanopore Technologies) sequencing generated 850,50 reads for DsR34 and 1,507,071 reads for DsR207, sufficient for 448 and 527 mean genome coverage and enabled closure of the genome with high confidence. Mutations in DsR34, and DsR207 mutants were identified by mapping against the *D. solani* IPO 2222 reference genome ^29^. In mutant DsR34 a mutation in the gene encoding secretion protein HlyD conferred ΦD5 resistance. In mutant DsR207 a mutation in the gene encoding elongation factor Tu (EF-Tu) was responsible for phage resistance (Table 1).

**Table 1.**
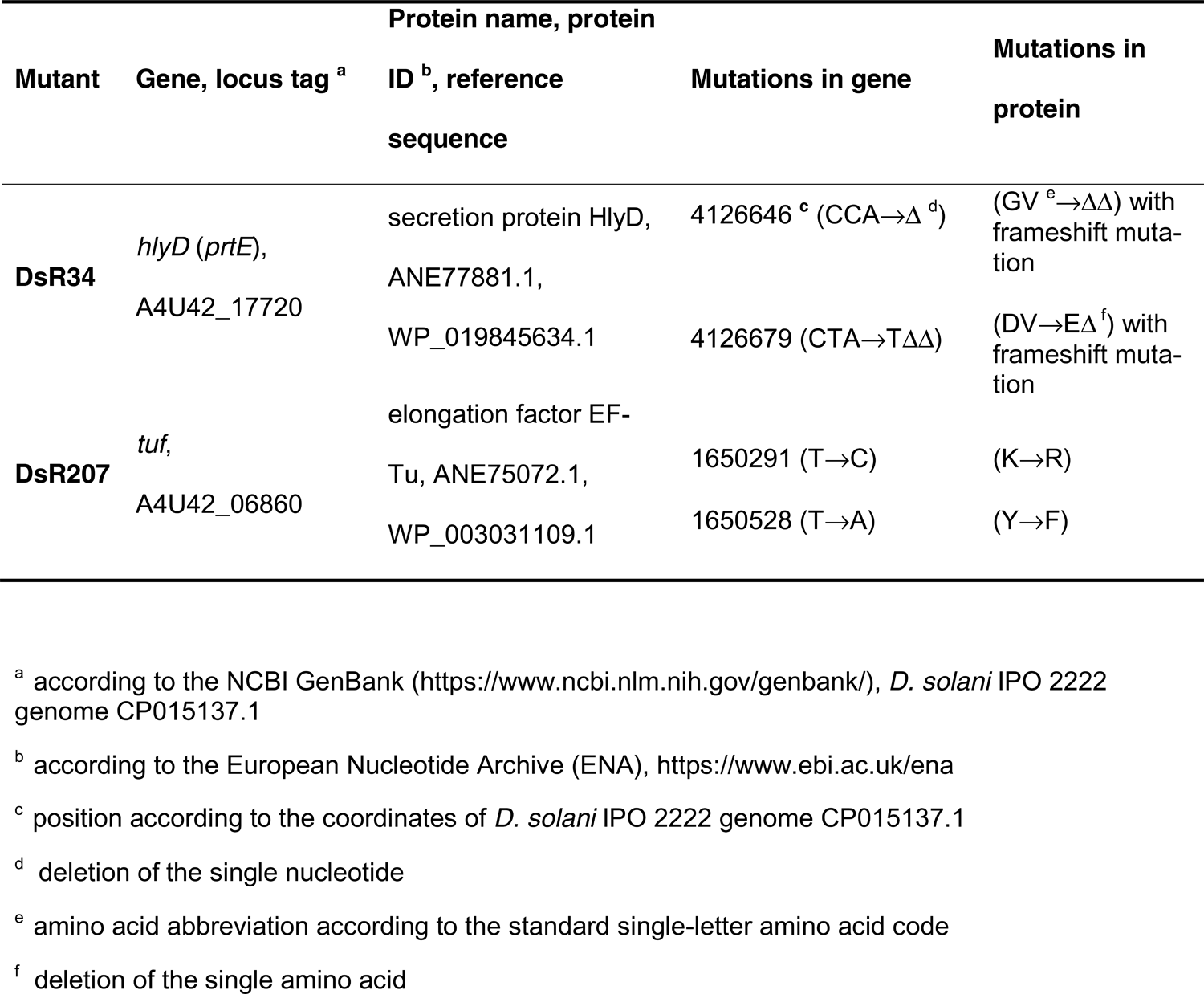
Genes mutated in the spontaneous phage-resistant *D. solani* mutants DsR34 and DsR207 found in this study

### Adsorption of ΦD5 to D. solani IPO 2222 wild-type strain and phage-resistant DsR34 and DsR207 mutants in vitro

The adsorption of ΦD5 to wild-type *D. solani* cells was fast. Within 5 minutes, nearly 90% of phage particles had absorbed to the wild-type strain and more than 95% had bound to this host by 20 minutes (Fig. 4). In contrast, the adsorption of the phage particles to cells of mutants DsR34 and DsR207 was significantly greatly reduced. Only between 1 and 25% of ΦD5 particles had bound to phage-resistant DsR34 and DsR207 cells within 20 minutes (Fig. 4).

**Figure 4.**
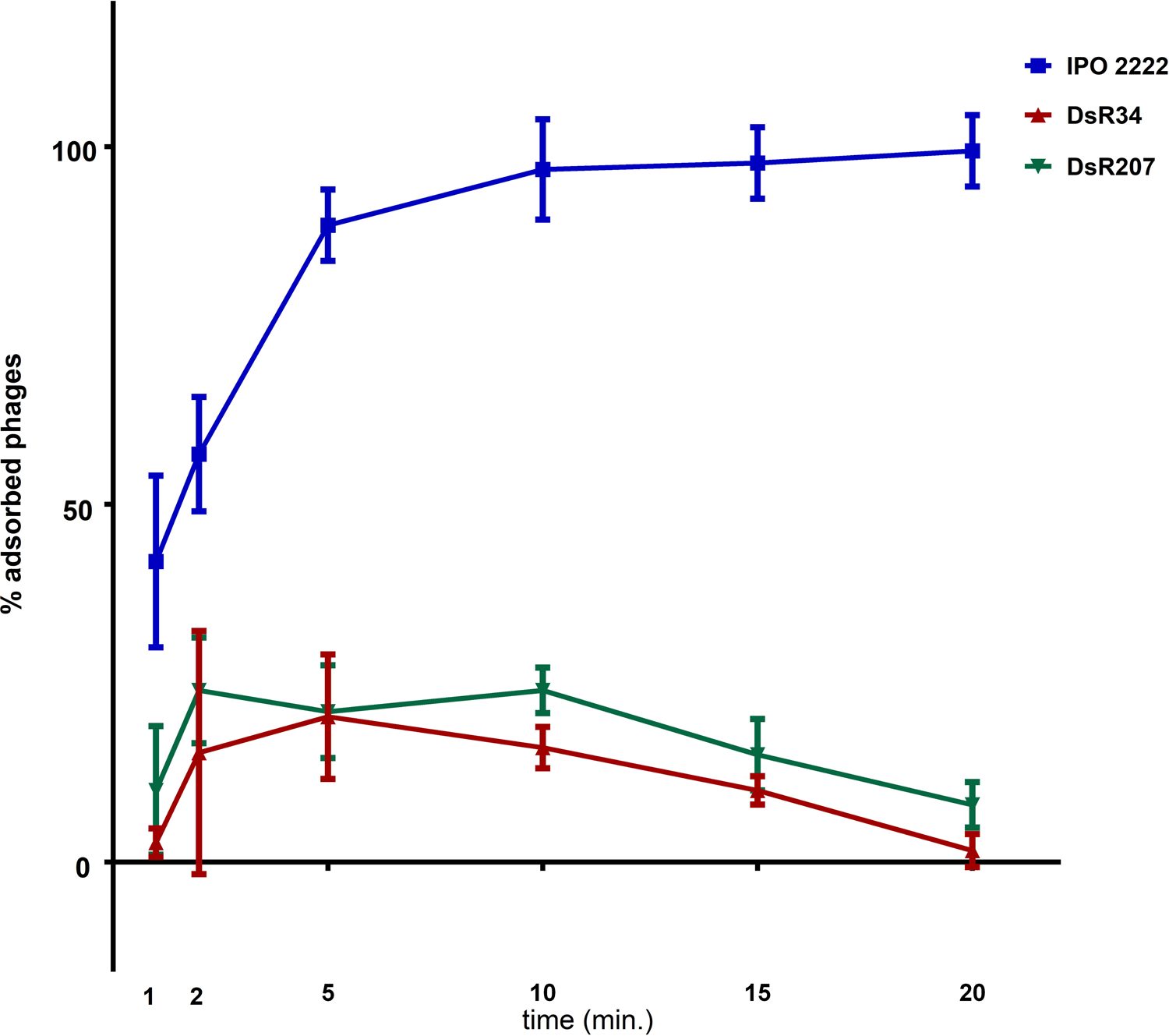
Adsorption of ΦD5 to cells of *D. solani* IPO 2222 wild-type strain and phage-resistant mutants Dsr34 and DsR207. A MOI of 0.01 of ΦD5 was used for the adsorption assay. The total time of the assay was 20 min. Phage adsorption was calculated as described previously ^42^: the percentage adsorption = (the average titer of unabsorbed phages per sample/average titer of phages in negative control) ×100. The averages and standard deviations (SD) of three independent repetitions per strain (WT or phage-resistant mutants) are shown.

### Phenotypes of phage-resistant DsR34 and DsR207 mutants

The phage-resistant mutants were analyzed for changes in their phenotypes that might be essential for their ecological fitness and virulence in natural and agricultural settings ^30–32^. There were no differences between the mutants and the wild-type strain in most phenotypes examined in this study. Specifically, phage-resistant mutants, like the wild-type strain, were capable of producing pectinases, forming cavities on the CVP medium, produced proteases, degraded carboxymethylcellulose, and polygalacturonic acid. They all were unable to produce siderophores or grow on medium supplemented with 5% NaCl. Likewise, no difference between WT *D. solani* and phage-resistant mutants DsR34 and DsR207 was observed in the production of secreted enzymes tested with API-ZYM assays. Furthermore, no differences in cell morphology or size diameter were observed in the phage-resistant mutants compared to the IPO 2222 strain by examination using transmission electron microscopy (TEM) (Fig. 5). Implementing atomic force microscopy (AFM) imaging did not reveal significant differences in the surface morphology of the analyzed *D. solani* phage-resistant cells with the scanning method applied. The bacteria were flagellated, and their measured length, width, and height were like the *D. solani* IPO 2222 wild-type strain (Fig. 6). The phage-resistant *D. solani* mutants exhibited comparable morphologies and colony size as that of the WT strain. None of the mutants differed significantly in their generation times in either rich (TSB) or minimal (M9 + glucose) media compared to the wild-type strain. The growth rate of the mutants tested at various temperatures (5, 8, 15, 28, and 37 °C) did not differ from that of the WT strain. The growth of the mutants at pH 4.0 and 10.0 was also similar to that of the wild-type strain. Phage-resistant mutants expressed equivalent susceptibility/resistance to all antibiotics tested as the WT strain. While the WT strain could move by swimming and swarming, both phage-resistant mutants were capable of swimming motility but not swarming motility. No significant differences were observed between the phage-resistant mutants and the wild-type IPO 2222 strain in most metabolic phenotypes examined. Both DsR34 and DsR207 mutants differed from the wild-type IPO 2222 strain in a total of only 9 features out of 316 such metabolic traits tested using BIOLOG phenotypic microarrays. All these divergent traits (except their resistance to 4% NaCl), were related to carbohydrate metabolism. The phage-resistant mutants both lost the ability to utilize D-cellobiose, D-turanose, and gentiobiose and became susceptible to 4% NaCl. Simultaneously, DsR34 and DsR207 mutants each gained the ability to use N-acetyl-β-D-mannosamine, L-glutamic acid, inulin, D-tagatose, and malonic acid for growth. Phage-resistant mutants DsR34 and DsR207 exhibited less rapid sedimentation associated with self-aggregation than the wild-type strain (Fig. 7). The SDS-PAGE patterns of the lipopolysaccharides purified from the ΦD5-resistant mutants DsR34 and DsR207 were indistinguishable from each other. However, compared to the LPS purified from the WT strain, these LPSs differed by the absence of one faint band of ca. 30 kDa (Fig. 8).

**Figure 5.**
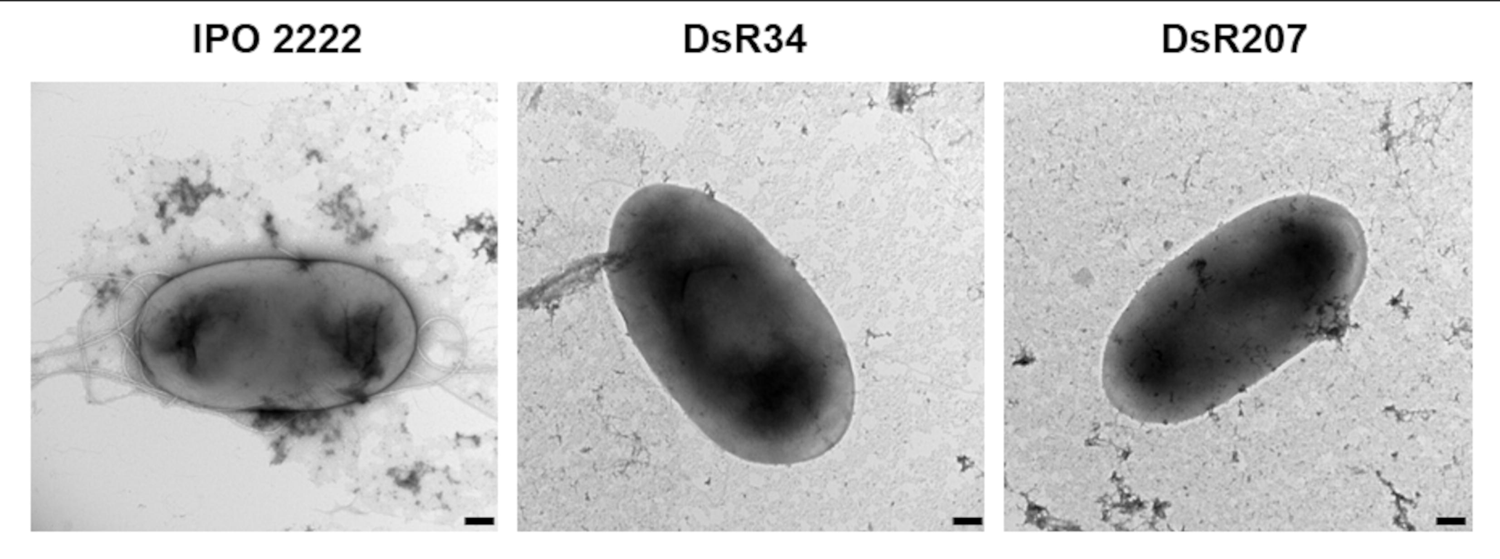
Morphology of *D. solani* IPO 2222 wild-type strain and phage-resistant DsR34 and DsR207 mutants visualized by transmission electron microscopy (TEM). TEM analyses were conducted on bacterial cells grown overnight in Tryptone Soya Broth (TSB) with shaking (200 rpm) at 28 °C. Photos were taken directly after the collection of the bacteria from liquid cultures. For TEM analysis, bacteria were adsorbed onto carbon-coated grids (GF Microsystems) stained with 1.5% uranyl acetate and directly examined with the electron microscope (Tecnai Spirit BioTWIN, FEI). At least ten images were taken per analyzed strain. The figure shows representative images. The bar represents 200 nm.

**Figure 6.**
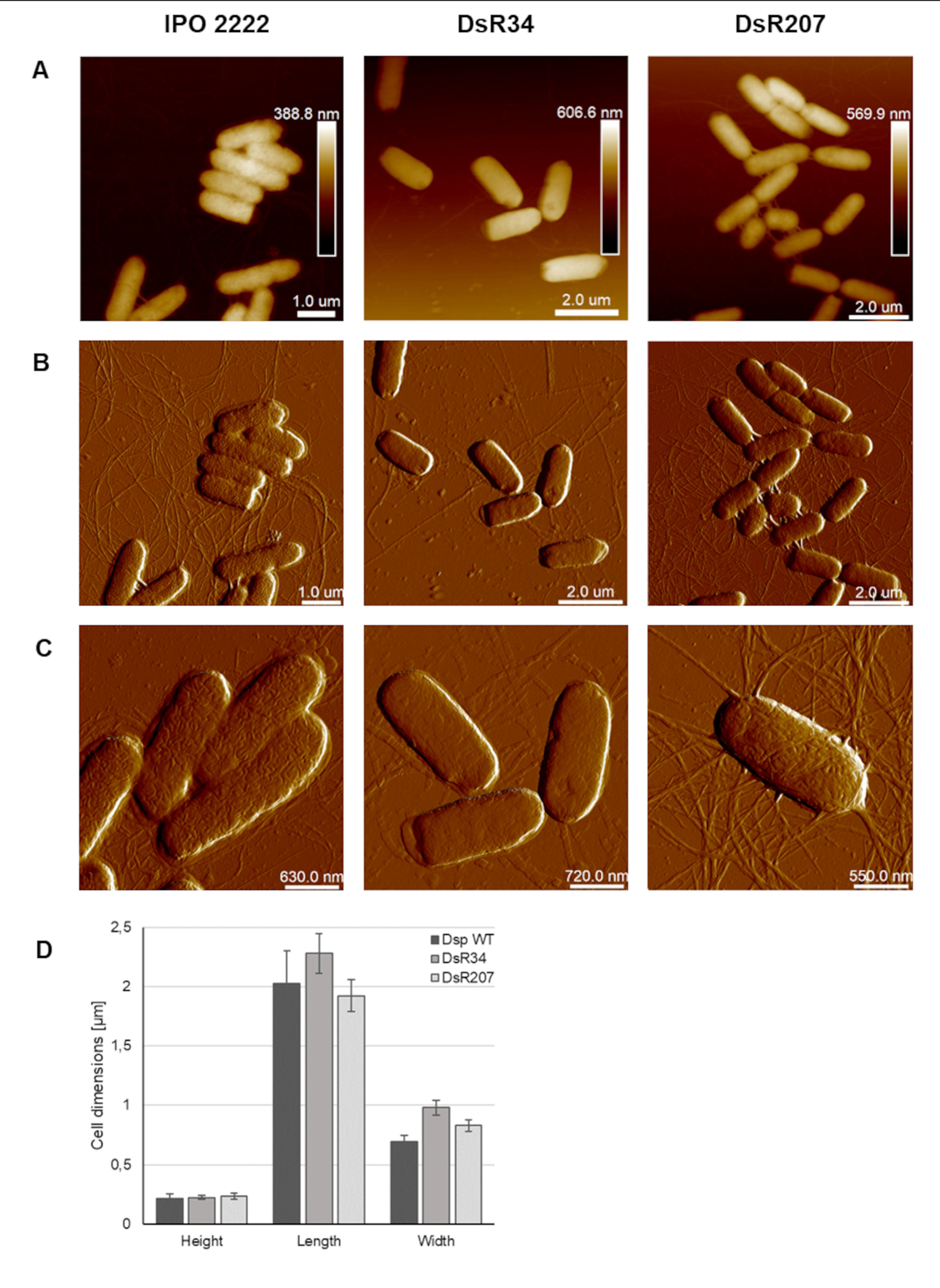
Morphology of *D. solani* IPO 2222 wild-type strain and phage-resistant DsR34 and DsR207 mutants visualized by atomic force microscopy (AFM). AFM analyses were conducted on bacterial cells grown overnight in M9 +0.4% glucose on a microscope glass slide with gentle shaking (60 rpm) at 28 °C. The cells were fixed with 2.5% glutaraldehyde for 2 h, washed, and dried in air. Cells were imaged using Bioscope Resolve (Bruker) in ScanAsyst (Peak Force Tapping) mode using ScanAsyst Air probe (f0 7.0 kHz, diameter <12 nm, k:0.4 N/m). **A** - Height and peak force error, **B** and **C** - Images of the analyzed cells. **D** - Dimensions of the bacteria were measured for N=12 to 20 cells, for their height, length, and width are shown as means with standard deviation (SD) for each strain.

**Figure 7.**
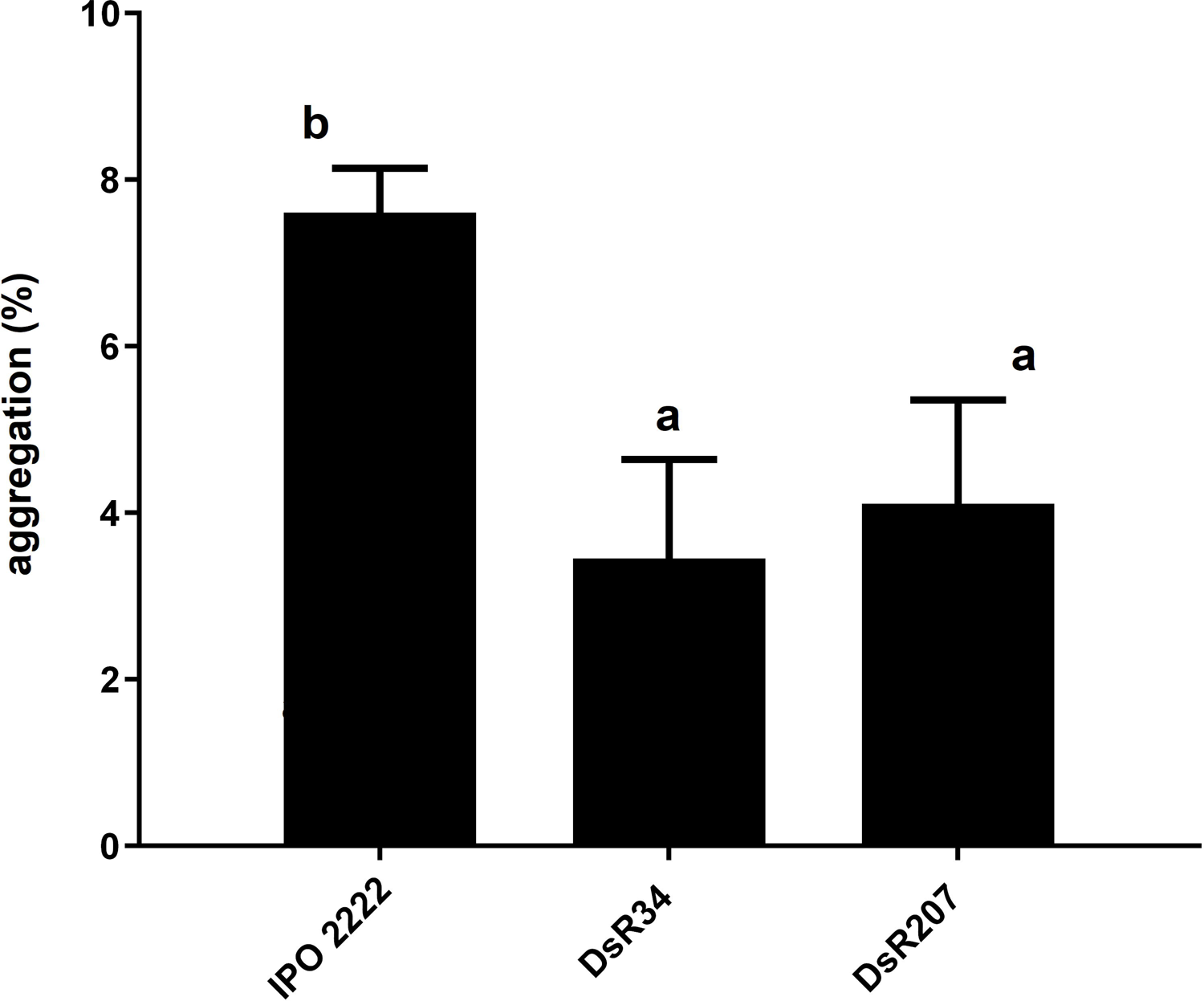
Self-aggregation of cells of *D. solani* IPO 2222 wild-type strain and phage-resistant DsR34 and DsR207 mutants as measured by turbidity of the bacterial suspension. The percentage of aggregation was quantified from the change in optical density (OD_600_) over 24 h. Percentage aggregation (sedimentation) was measured as follows: %A = 1-(OD_600_ 24h/OD_600_ 0h), where: %A— the percentage of aggregation (sedimentation), OD_600_ 0h—OD of bacterial culture at time 0 h, OD_600_ 24h— OD of bacterial culture at time 24 h. Results were considered significant at p = 0.05, and pairwise differences were obtained using the *t*-test. The means not sharing the same letters above each bar vary. Error bars represent standard deviation (SD).

**Figure 8.**
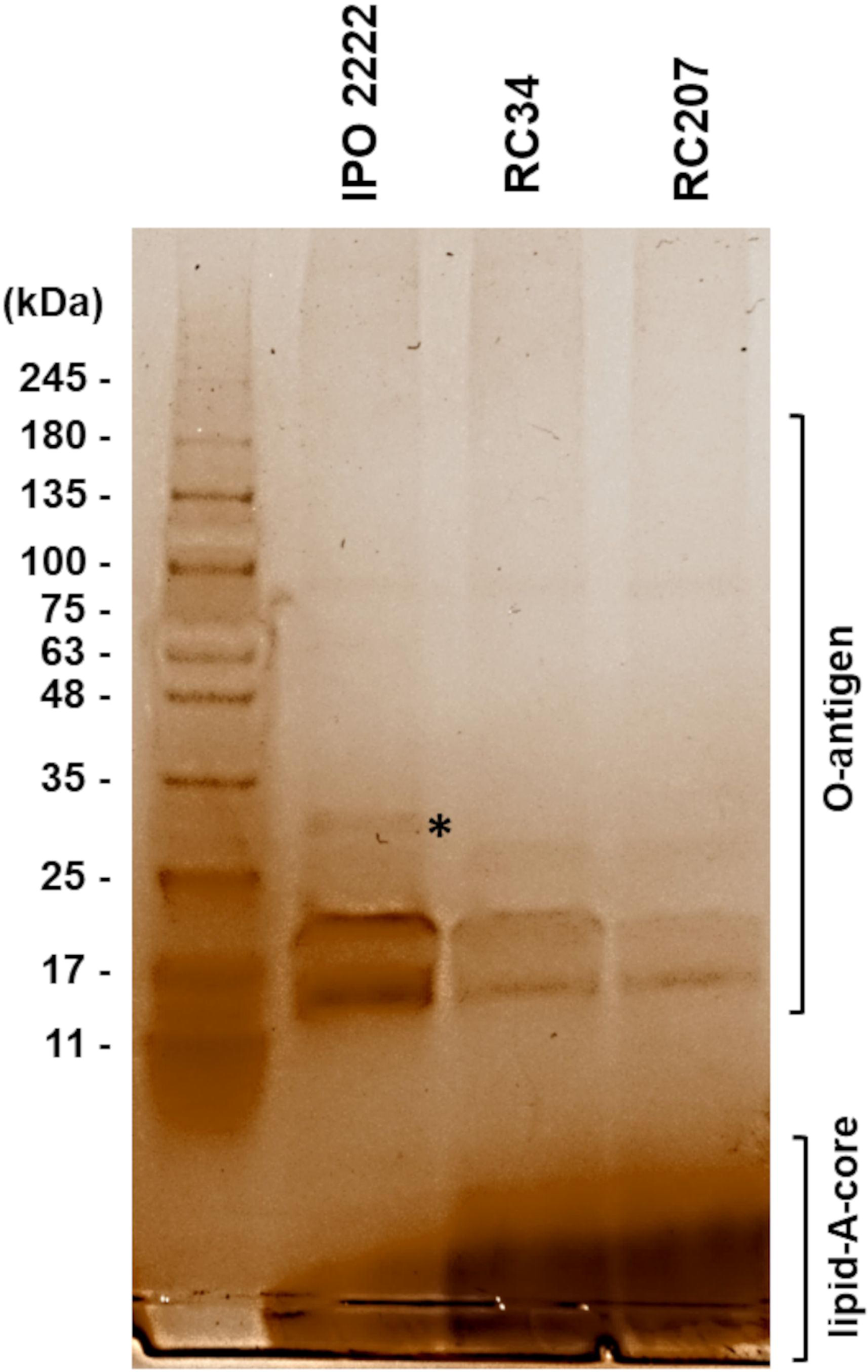
Characterization of lipopolysaccharide (LPS) from *D. solani* IPO 2222 wild-type strain and phage-resistant DsR34 and DsR207 mutants. SDS-PAGE was done using a gradient (4–20%) poly-acrylamide gel, and the LPS components were visualized by silver staining ^83^. The kDa marker (11– 245 kDa, Perfect Tricolor Protein Ladder, EURx, Poland) is shown in the first lane. **\*** - marks band present in the wild-type strain but absent in phage-resistant DsR34 and DsR207 mutants.

### Proteomics of the phage-resistant mutants grown in rich (TSB) and minimal (M9+glucose) media

To gain more insights into the characteristics of the spontaneous ΦD5-resistant mutants, the proteomes of the DsR34, DsR207, and wild-type strain grown in rich (TSB) and minimal (M9+glucose) media were compared using Sequential Window Acquisition of All Theoretical Mass Spectra (SWATH-MS) analysis. Differential protein expression was determined by comparing the relative abundance of proteins in phage-resistant mutants and the wild-type strain. Fold changes were considered significant with the q-value (adjusted p-value) <0.05.

Spontaneous phage-resistance affected DsR34 and DsR207 proteomes in both media tested. Among the analyzed strains, SWATH-MS identified a total of 492 *D. solani* proteins in the rich medium and 672 proteins in the minimal medium. In the rich medium, the core proteome of the WT, DsR34, and DsR207 consisted of 385 proteins, whereas 13, 15, and 24 unique proteins were significantly differentially expressed in cells of the WT, DsR34, and DsR207, respectively (Fig. 9A). Similarly, in the minimal medium (M9+glucose), the proteome shared between *D. solani* wild-type strain and phage-resistant mutants comprised 491 proteins, whereas 14, 48, and 30 proteins were uniquely differentially expressed in WT, DsR34, and DsR207, respectively (Fig. 9C).

**Figure 9.**
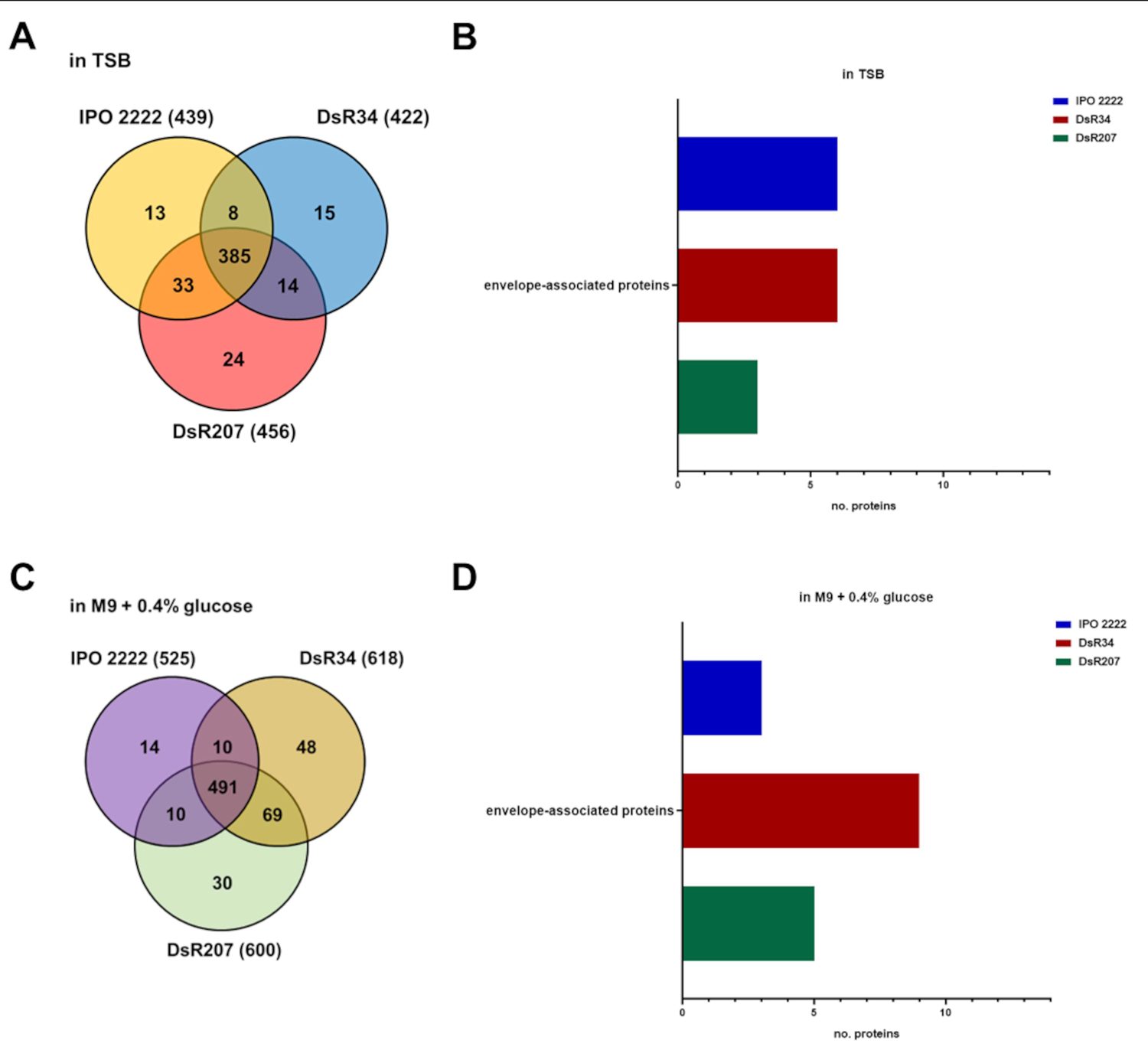
Proteomes of phage-resistant *D. solani* mutants DsR34 and Dsr207. **A, C** – Venn plot illustrating protein identification overlaps between the wild-type IPO 2222 strain and ΦD5-resistant mutants DsR34 and DsR207, **B, D** - Significantly differentially abundant proteins related to the envelope biogenesis, metabolism, and regulation in phage-resistant mutants DsR34 and DsR207 and the wild-type IPO 2222.

The unique proteins (=significantly differentially expressed) were grouped into ten categories according to their primary functions, *viz*. (1) protein metabolism, (2) carbohydrate metabolism, (3) lipid metabolism, (4) DNA metabolism, (5) transport and signaling, (6) stress and redox processes, (7) metal ion homeostasis, (8) motility and chemotaxis, and (9) co-factors metabolism and (10) others (miscellaneous proteins with unknown function and/or localization) (Supplementary Fig. 1). As expected, the number of proteins in a particular category differed between the analyzed strains and the type of medium tested (Fig. 9A & C). However, among the tested strains, most of the differently expressed proteins were represented by the category protein metabolism, and category of others (proteins with unknown function and localization) in both media tested. Proteins related to DNA metabolism, transport and signaling, metal ion homeostasis, and motility and chemotaxis were not more abundant in the WT strain in the rich medium (TSB) relative to that of the phage mutants.

Likewise, proteins related to lipid metabolism, metal ion homeostasis and, motility and chemotaxis were not differentially abundant in the WT strain relative to the mutants in a minimal medium (M9+glucose). The proteins upregulated in DsR34 in the rich medium were linked with carbohydrate metabolism, transport and signaling, motility, and chemotaxis. Contrary, in the minimal medium, proteins associated with protein metabolism, DNA metabolism, transport and signaling and stress and redox processes were upregulated in this mutant. In the case of the DsR207 mutant, in the rich medium, proteins belonging to two categories were significantly upregulated: those involved in protein metabolism and proteins in the category ‘others’ with unknown functions. In mutant DsR207, proteins associated with carbohydrate and lipid metabolism were upregulated in the minimal medium compared to the other strains (Supplementary Fig. 1).

Given that the two phage-resistant mutants might be expected to have alternations in their cell surface that resulted in the decreased ΦD5 adsorption, we compared the proteomes of DsR34 and DsR207 mutants and WT strain for differentially expressed proteins associated with bacterial cell surfaces (=envelope, transmembrane transport, and signaling) (Table 2). Such proteins constituted 40 and 19% of the total unique proteins expressed in rich and minimal medium, respectively for mutant DsR34. In rich media the fraction of envelope-associated proteins among all unique proteins was 12.5%, whereas, in minimal medium, this fraction was 17% for mutant DsR207 (Fig. 9B & D).

**Table 2.**
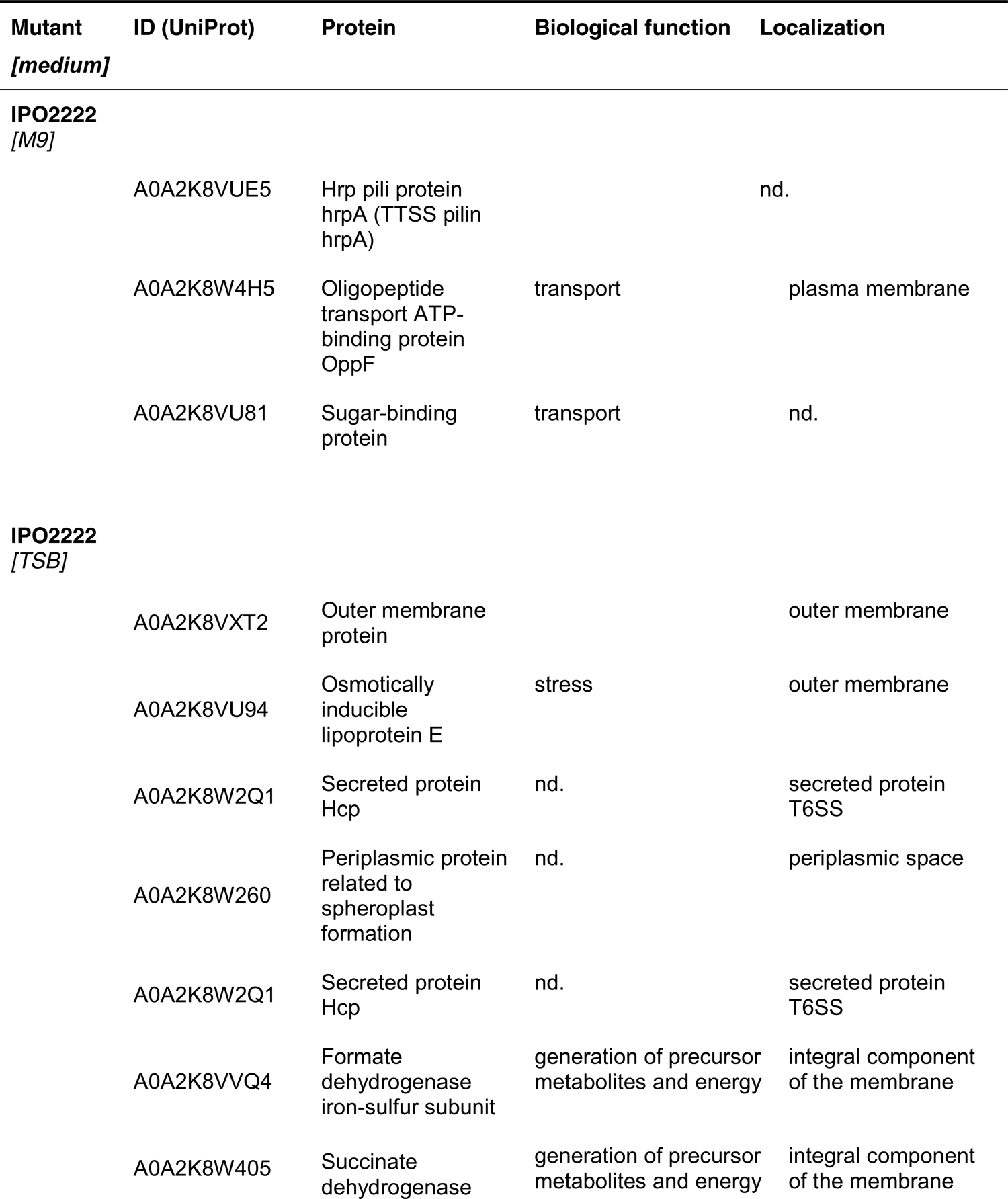
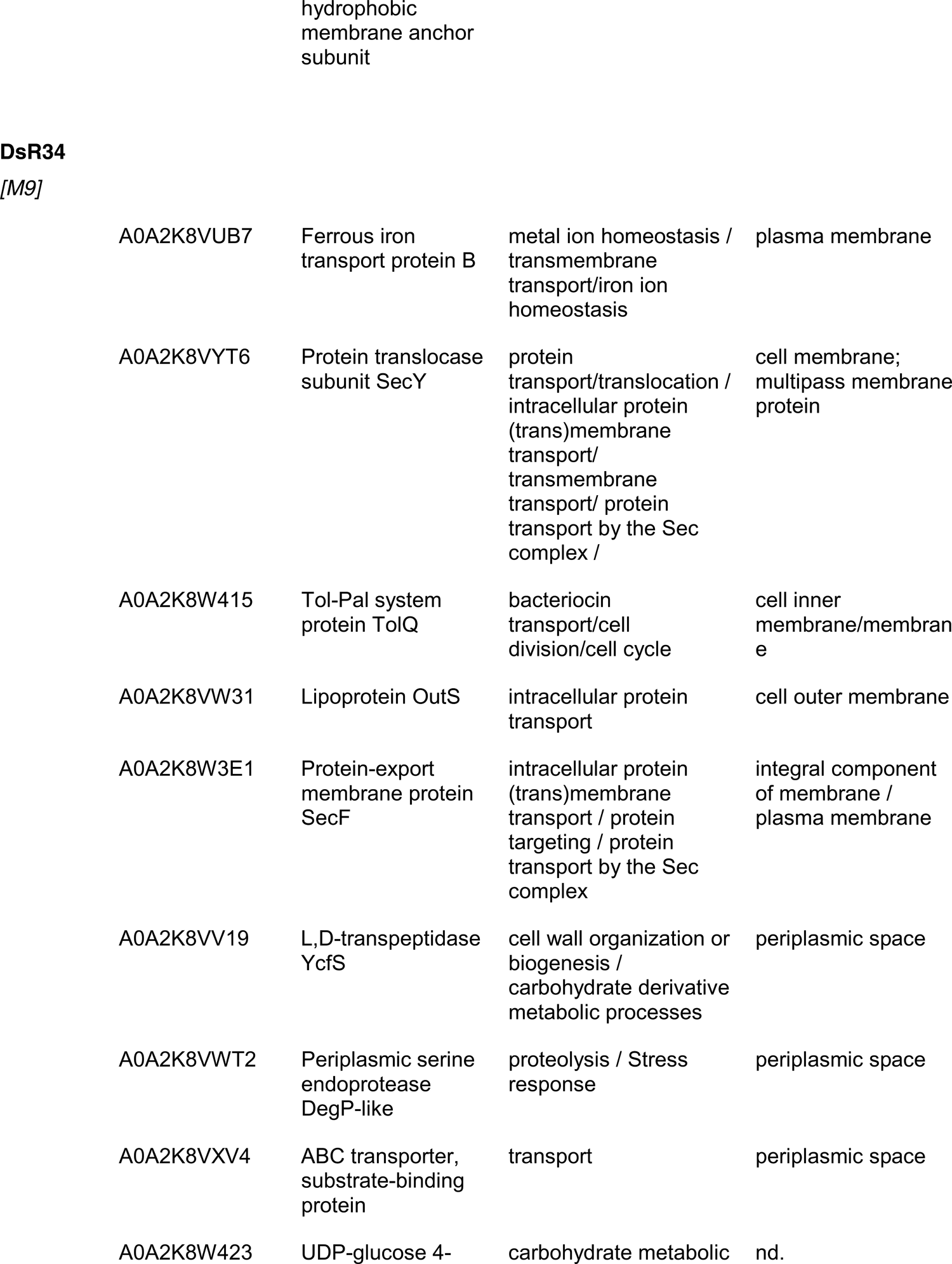
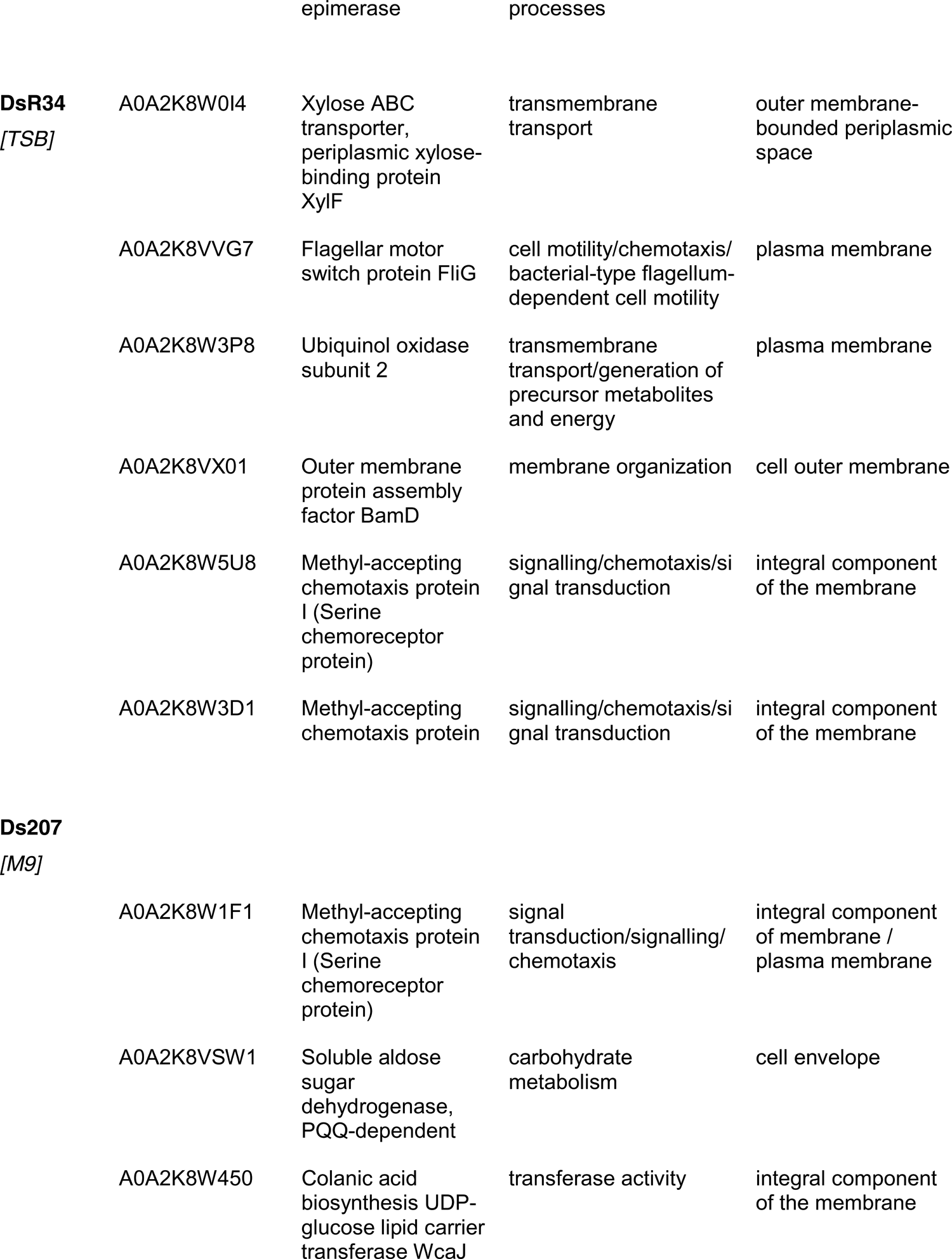
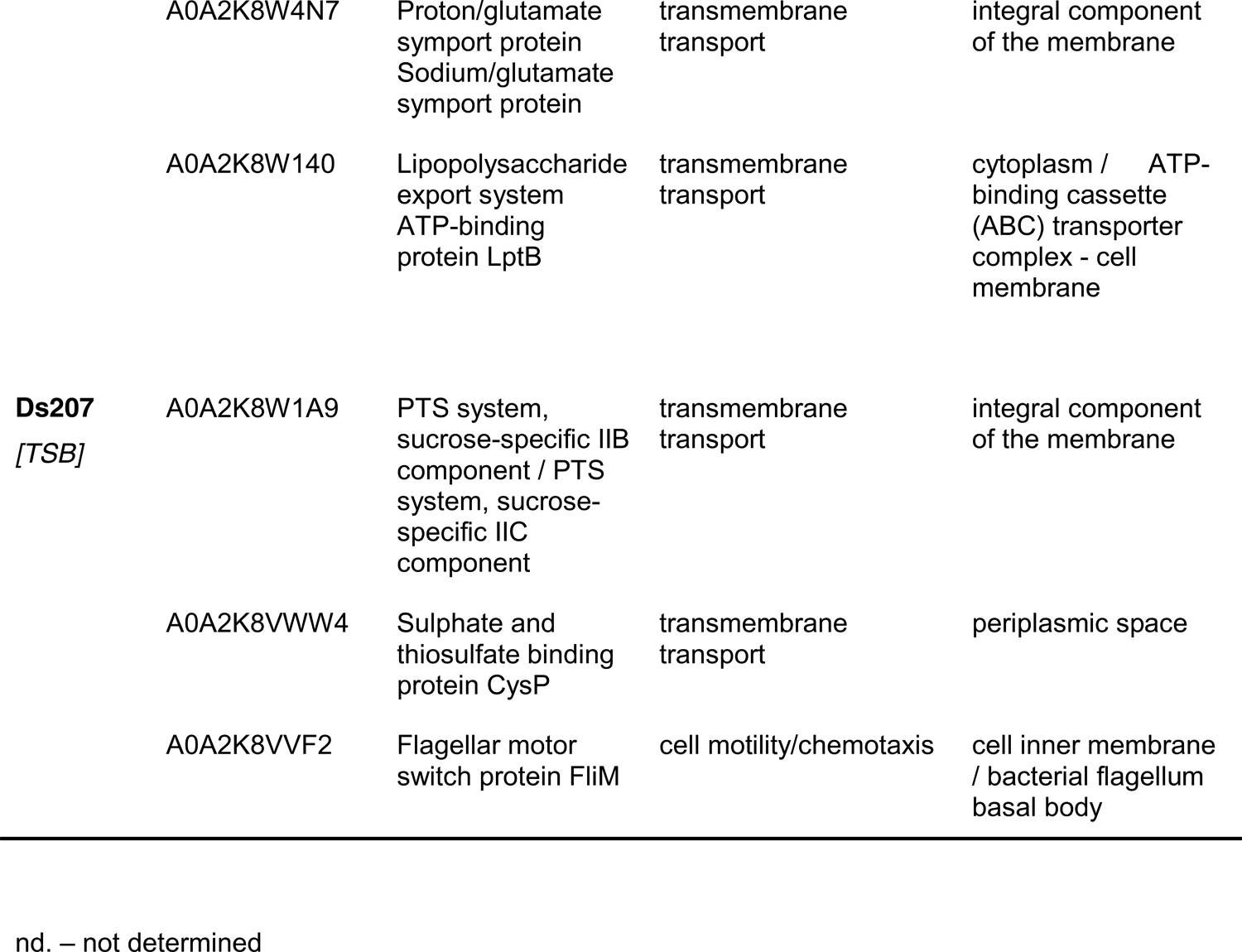
Unique proteins of phage-resistant DsR34 and DsR207 mutants associated with the bacterial envelope, transmembrane transport, and signaling of *D. solani*

### Effect of selected surfactants on the viability of phage-resistant D. solani mutants

No differences were found between the phage-resistant mutants DsR34 and DsR207 and *D. solani* IPO 2222 wild-type strain in response to Triton X-100, Tween 20, Tween 80, Poloxamer 407, and N-lauroyl sarcosine. In contrast, both phage-resistant mutants exhibited reduced growth in the presence of 0.1 % EDTA compared with that of the WT strain. On average, the cell concentrations DsR34 and DsR207 after 16 h incubation in the presence of 0.1 % EDTA were only about half that of the WT strain (Fig. 10).

**Figure 10.**
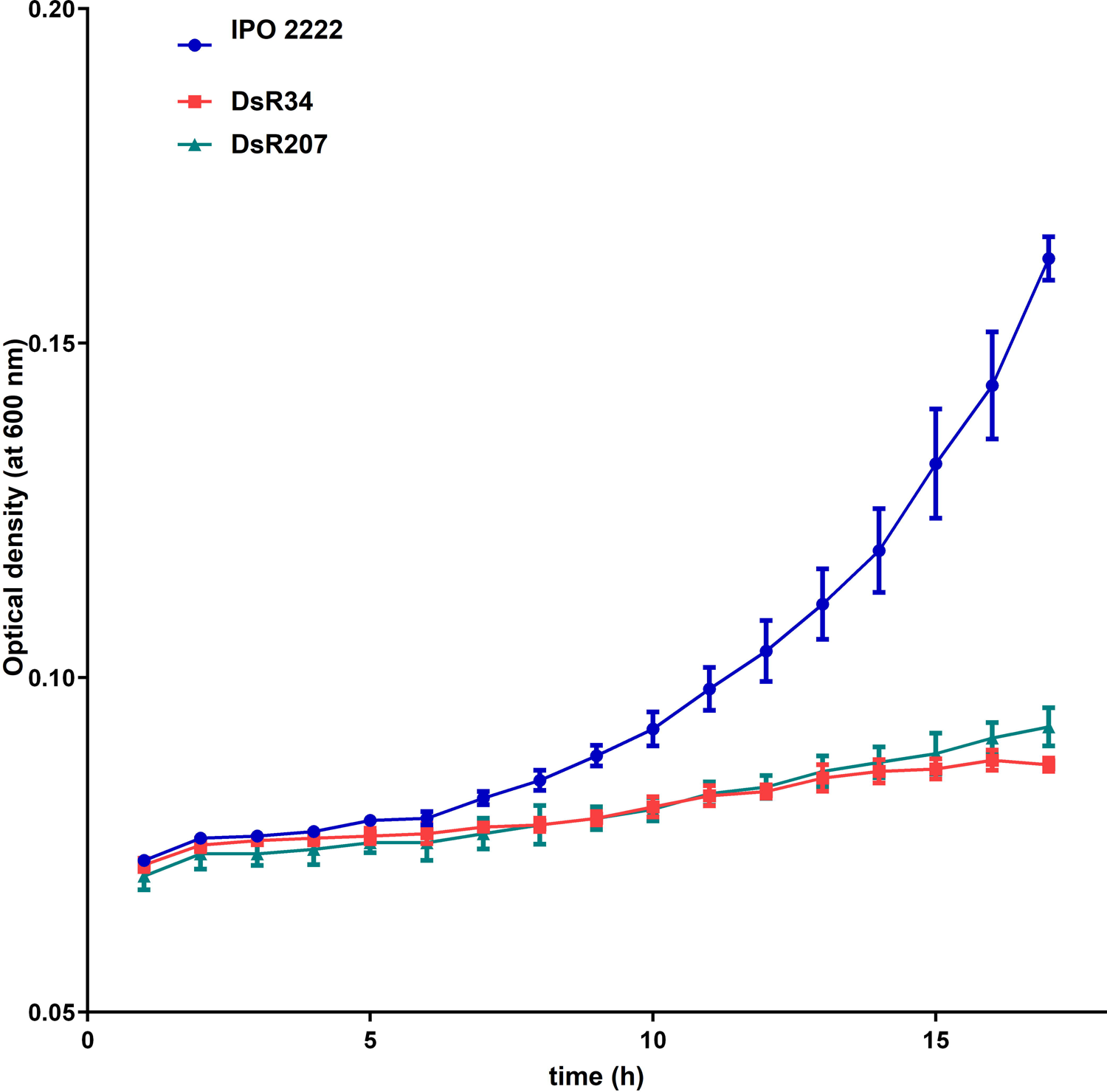
Growth of *D. solani* IPO 2222 wild-type strain and phage-resistant DsR34 and DsR207 mutants in Tryptone Soya Broth (TSB) supplemented with 0.1% EDTA. The experiment was done using two biological replicates containing two technical replicates each (n=4). The results were averaged for presentation. The bars show the standard deviation (SD).

## Discussion

Our understanding of the mechanisms underlying interactions of Soft Rot *Pectobacteriaceae* bacteria with lytic bacteriophages remains limited ^16, 26^. This study explored whether there is a direct link between spontaneous phage resistance and virulence of plant pathogenic *D. solani* strain IPO 2222. By focusing on virulence and interrogating the spontaneous phage-resistant mutants in plant-related environments, we wanted to assess whether resistance to phage infections impacts the ecological success of the pathogen, defined as the ability to survive in the plant surrounding and efficiently colonize and cause symptoms in plant hosts.

Although most ΦD5-resistant *D. solani* mutants found in this study were unaffected in virulence (>99%), we identified two mutants, DsR34 and DsR207, that exhibited a significant reduction in their ability to macerate potato tuber tissues. It was perhaps not a surprise that most phage-resistant *D. solani* mutants retained virulence. It is of utmost importance for a pathogen to remain virulent ^33, 34^, as losing the ability to infect the host heavily impacts its ability to generate large population sizes thus its survival and ecological success ^35^. On the other hand, in complex environments such as the surface and interior of plants, the evolution of resistance to phage infections is likely to be very costly. Phage selective pressure is expected to be a strong driver of spontaneous mutations ^36^. In contrast, a direct link between spontaneous resistance to viral infections and reduced virulence has already been shown for members of several bacterial species ^37^, including plant pathogenic bacteria, but to our knowledge, not for *D. solani*. For example, spontaneous resistance to bacteriophage X2 by the in-plant pathogen *Xanthomonas oryzae* pv. *oryzae* resulted in reduced virulence ^38^. Reduced virulence was also observed in the case of spontaneous resistance of *Pseudomonas syringae* pv. *porri* to phages KIL3b and KIL5 ^39^. It suggests a direct tradeoff cost for phage-resistant mutants in a plant environment. Likewise, recent studies on human pathogens *Staphylococcus aureus* ^40^ and *Serratia marcescens* ^38^ suggested that selective pressure conferred by the presence of lytic bacteriophages results in the appearance of spontaneous phage-resistant mutants having a lower virulence. Such observations highlight the global link between viral resistance and the ability to cause infections by bacterial pathogens ^41^.

Gram-negative bacteria have developed several strategies to cope with viral infections ^7^. Among different mechanisms of phage evasion, modification of the bacterial cell surface that prevents phage attachment has been the most often reported ^10^. In this study, the adsorption of ΦD5 to *D. solani* mutants DsR34 and DsR207 was greatly reduced. This indicates that probably the preferred mechanism of spontaneous resistance against phage ΦD5 in *D. solani* IPO 2222 is the modification of the cell surface in a way to be unsuitable for a phage attachment (*i.e.*, adsorption inhibition) ^11^.

The sequencing of the DsR34 and DsR207 genomes showed that spontaneous ΦD5 resistance was caused by mutations in two distinct loci not connected so far with phage resistance in SRP bacteria ^42^ *viz*. in the gene coding for secretion protein HlyD (mutant DsR34) and in the gene encoding an elongation factor Tu (EF-Tu) (mutant DsR207). HlyD secretion protein is a conserved trimeric transmembrane fusion protein and a part of the type I secretion system (T1SS) in Gram-negative bacteria. HlyD is involved in the secretion of α-hemolysin in *Escherichia coli* ^43^. In *Dickeya* spp., a HlyD homolog, protein PrtE takes part in protease secretion ^44^, contributing to virulence *in planta* ^45, 46^. A Δ*prtE* mutant exhibited decreased virulence due to its lack of various secreted proteases during infection ^47^. Nevertheless, in our study, the DsR34 mutant maintains the ability to produce proteases. PrtE/HlyD has not yet been previously connected with phage-host interactions, including interactions of Soft Rot *Pectobacteriaceae* members with their bacteriophages.

The other mutated protein, elongation factor Tu (EF-Tu), is a G protein catalyzing binding of aminoacyl-tRNA to ribosomes ^48^. As such, it has an important role in translation. In addition to its primary function in translation, EF-Tu has the ability to execute diverse tasks on the surface of bacterial cells, including interaction with membrane receptors and the extracellular matrix ^48^. In addition, EF-Tu is a recognized pathogen-associated molecular pattern (PAMP) that is recognized by plants ^49, 50^. In this study, DsR207 mutant has two missense mutations in the EF-Tu gene. It can be speculated that these two mutations may altered the protein in a way that the mutated version is better recognized as a PAMP by the plant than the wild-type EF-Tu ^51^, resulting in a strong immune response, thus reducing the DSR207 virulence. It was reported that EF-Tu is important in the interaction of several plant pathogens, including *P. carotovorum*, *Ralstonia solanacearum,* and *Agrobacterium tumefaciens,* with their host plants acting as an elicitor of the plant innate immune response ^52^. Although no reports clearly link EF-Tu with phage-SRP bacteria interaction, it is known that *Escherichia coli* EF-Tu participates in the bacteriophage exclusion system (altruistic suicide of infected cells upon infection) ^53^, preventing propagation of T4 phage in the environment. It remains unclear whether *Dickeya* spp. EF-Tu may work in a similar way as in *E. coli*.

Spontaneous resistance of DsR34 and DsR207 against ΦD5 had pleiotropic phenotypic consequences. Both mutants identified in this study retain several phenotypic alternations that differentiate them from the WT strain, which may decrease their fitness *in planta*. Both mutants were unable to swarm, were significantly more susceptible to EDTA, and expressed less rapid cell sedimentation/aggregation compared to the wild-type IPO 2222 strain. Swarming is known to be one of the vital features needed by *D. solani* to colonize and establish an infection in plants ^54^. It is therefore not a surprise that inability to swarm observed in phage-resistant mutants may result in decreased virulence of the DsR34 and DsR207 mutants *in planta* ^35^. Similarly, EDTA is known to increase the permeability of the capsule in Gram-negative bacteria, making them more susceptible to environmental stresses ^55^. Likewise, differences in self-aggregation of phage-resistant mutants compared to the wild-type strain suggested changes in the surface features of *D. solani* cells ^56^.

Likewise, spontaneous mutations conferring phage resistance directly affected the proteomes of DsR34 and DsR207 mutants, including the abundance of the proteins associated with cell envelope in *D. solani* strain IPO 2222. Although the two mutations found in this study have not yet been linked to the resistance of SRP or other Gram-negative bacteria to viral infections ^7^, they affected the expression of various proteins associated with transmembrane transport, cell wall organization, and metabolism of envelope-associated carbohydrates. Such a phenotype directly links phage resistance with the status of the bacterial envelope, a common feature of phage resistance. Consistent with the conjecture, analysis of the LPS derived from both DsR34 and DsR207 revealed altered LPS. Such a phenotype may explain the decreased attachment of phages to the cells of these mutants. In our earlier work, which analyzed the interaction of *P. parmentieri* with lytic phage ΦA38, we discovered that this phage required native LPS to infect its host ^57^. Likewise, we also showed previously that mutations targeting the LPS cluster in *D. solani* ^58^ resulted, among other phenotypes, in resistance to bacteriophage infections ^42^. It seems that modifications of LPS and capsule to prevent viral infections frequently occur in bacteria belonging to Soft Rot *Pectobacteriaceae*, as similar observations were made for other members of this group, including *P. atrosepticum*, *P. carotovorum*, and *P. brasiliense*^59–61^. The apparent differences in LPS structure, elevated EDTA susceptibility, and less rapid self-aggregation further strengthened the hypothesis that alternations of bacterial surface properties occurred in phage-resistant mutants DsR34 and DsR207 ^62^. It is not clear yet how the mutations in HlyD may result in alternations in *D. solani* surface properties, however it has been already shown that HlyD interacts with the LPS of *D. solani* ^45^. Therefore, it cannot be excluded that mutations of HlyD may result in its incorrect folding, insertion and localization of protein in the periplasm and by these may also affect the LPS structure ^45^. Another possibility is that mutations found in HlyD may cause misfolding of the secreted proteins and therefore contributed to the decreased virulence of DsR34 mutant. It was shown that point mutations of HlyD impacted the folding of the HlyA protein and its translocation in *E. coli* ^63^. Similar situation may take place in *D. solani*. The other mutated protein, EF-Tu, was reported to be a part of bacterial cytoskeleton, required to maintain the proper shape of the bacterial cell ^64^. EF-Tu localizes underneath the cell membrane of *E. coli* and *Caulobacter crescentus*^65^. In *Bacillus subtilis*, EF-Tu affects cell surface and the decrease in the concentration of EF-Tu leads to defects in cell surface morphology ^66^. The interaction of EF-Tu and cytoskeleton elements seems to by an universal mechanism by which the prokaryotic cells sustain their morphological features ^67^.

To support the role of *D. solani* surface alterations in phage avoidance, we used microscopic techniques to compare the surface of the wild-type strain and the spontaneous ΦD5-resistant mutants DsR34 and DsR207. Phage-resistant mutants were overall indistinguishable from the WT strain in the colony and cell morphology. To our surprise, also more detailed analyses done with TEM and AFM did not reveal any apparent alterations in the cell surface of the ΦD5-resistant mutants that could easily explain the observed phenotypes, including the lowered virulence *in planta*. Furthermore, the DsR34 and DsR207 cells possessed the same surface topography, diameter, and size as the wild-type IPO 2222. It is likely however, that the changes in cell surface LPS and proteins composition would have led to very modest changes in physical appearance of the envelope surface, with such changes being undetectable by even AFM.

The two phage-resistant mutants exhibited reduced virulence in detached chicory leaf assay as well as reduced colonization and symptom expression in potato plants growing in potting soil. Such results indicate that both mutants were heavily compromised in their ability to invade and multiply within different hosts ^51^. Their decreased virulence of the phage-resistant mutants after introduction into chicory leaves and in potato plants after soil inoculation was not due to intrinsic reductions in growth rate; under *in vitro* conditions, the mutants and the WT strain had comparable generation times under a number of conditions, including when grown at different temperatures, pHs, and in the presence of various carbon sources. It is clear, therefore, that the observed ΦD5 resistance strongly impacted both the multiplication of *D. solani in planta* and thus its ability to generate secondary inoculum for persistence and infection of other plants. For this reason, inhibition of phage adsorption by *D. solani* to avoid infections will likely reduce its ecological fitness in many cases. Such an observation may have practical importance, since implementing phage therapy against such an agricultural pathogen may result not only in the killing of the pathogen but also in the selection of less virulent phage-resistant variants that would reduce the future risk of infection by that species ^68^.

## Materials and Methods

### Bacteriophages, bacterial strains, and growth conditions

The lytic bacteriophage vB_Dsol_D5 (ΦD5), described previously ^69–71^, was grown on its host, *D. solani* strain IPO 2222 ^20^, and quantified as described earlier ^69^. A stock of phage particles (ca. 10^8^ – 10^9^ PFU (plaque-forming units) mL^-1^) in TSB (tryptone soya broth, Oxoid) or quarter-strength (1/4) Ringer’s buffer (Merck)) was used in this study. The wild type (WT) *D. solani* was cultivated for 24-48 h at 28 °C on tryptone soya agar (TSA, Oxoid), in tryptone soya broth (TSB, Oxoid) or M9 medium (MP Biomedicals) supplemented with 0.4 % glucose (final concentration 0.4%). To solidify the media, bacteriological agar (Oxoid) (15 g L^-1^) was added when needed. Spontaneous ΦD5-resistant *D. solani* mutants were cultivated under the same conditions and using the same growth media as the WT strain. To prevent fungal growth, the growth medium was supplemented with cycloheximide (Sigma-Aldrich, final concentration: 200 μg mL^-1^).

### Recovery of spontaneous phage-resistant mutants of D. solani

The recovery of ΦD5-resistant mutants of *D. solani* was made as previously described ^72^. Briefly, the overnight culture of the WT strain grown in TSB (rich medium) or M9+0.4% glucose (minimal medium) was diluted 50 times in the same fresh medium. The diluted bacterial culture was grown at 28 °C with shaking (120 rpm) to achieve an optical density (at 600 nm = OD_600_) of ca. 0.5. Cultures were then spiked with ΦD5 particles suspended in sterile TSB to reach the final multiplicity of infection (MOI) of 0.01 and incubated for the next 72 - 96 h under the same conditions. Bacteria surviving phage infections were purified into individual colonies by at least four passages on TSA agar plates ^73^ and collected for further studies. Verifying ΦD5 resistance of the obtained *D. solani* mutants was done as described previously ^42^. *D. solani* IPO 2222 mutants with confirmed ΦD5 resistance were selected for further studies.

### Confirmation of the identity of spontaneous phage-resistant mutants by Enterobacterial Repetitive Intergenic Consensus (ERIC) PCR fingerprinting

ERIC-PCR was done on selected spontaneous ΦD5-resistant mutants to confirm their identity as *D. solani* IPO 2222, as previously described ^74^. Total bacterial genomic DNA was purified using Qiagen Genomic DNA Purification Kits (Qiagen) using the protocol provided by the manufacturer. The DNA concentration was adjusted with sterile demineralized water to 100 ng µL^-1^. Primer sequences corresponding to ERIC-1R (ERICIR (5’-ATGTAAGCTCCTGGGGATTCAC-3’) and ERIC2 (5-AAGTAAGTGACTGGGGTGAGCG-3 were used ^74^. The ERIC-PCR was performed with 25-μL volumes containing Taq DNA polymerase buffer 1X (Roche), 200 mM dNTP (Sigma-Aldrich), 0.4 mM of each primer, and 2 U of Expand High Fidelity Taq DNA polymerase (Roche) and 40 PCR cycles. Amplified DNA was analyzed by electrophoresis in 1.5% agarose gel in 0.5 x TBE buffer stained with 5 mg mL^-1^ of ethidium bromide (Sigma-Aldrich). Gels were developed for 6-7 h at 100 V at room temperature (approx. 20 – 24 °C). A 1 kb DNA ladder (Promega) was used as a size marker. The ERIC-PCR DNA patterns obtained for spontaneous phage-resistant mutants and WT strain were compared.

### Assessment of virulence of spontaneous phage-resistant mutants

The virulence of spontaneous phage-resistant mutants was tested initially using a whole potato tuber assay ^42^. Briefly, potato tubers (5 replicate tubers per bacterial strain) of cv. Bryza, purchased locally in Gdansk, Poland, were used. They were chosen for their similar size (diameter of ca. 5–6 cm and weight of ca. 50-70 g) ^42^ and inoculated with a given strain and evaluated for disease symptoms ^31, 75^. Bacterial suspensions (ca. 10^8^ CFU mL^−1^) (100 μl per strain/inoculation site) were delivered to the potato tuber by stab inoculation into the tuber pith with a 200 μl pipette tip. Inoculated tubers were kept in humid boxes (ca. 90% relative humidity) at 28 °C for 72 h to promote the expression of disease symptoms. Positive control tubers were inoculated with the WT *D. solani* strain, and the negative control tubers were inoculated with sterile demineralized water. The experiment was repeated once. Phage-resistant *D. solani* mutants demonstrating reduced ability to macerate potato tuber tissues were selected for further analysis.

### Sequencing of the genomes of selected ΦD5-resistant bacterial mutants, comparison with the wild-type genome, and identification of nucleotide and amino acid polymorphisms

DNA of selected phage-resistant mutants was isolated using a Wizard Genomic DNA purification kit (Promega Corp.) according to the guidelines provided by the manufacturer. Genome sequencing was done using second and third-generation of next-generation sequencing techniques ^76^. For short-read sequencing, samples were quantified and diluted according to service provider specifications and sent offsite for processing. Long-reads (Oxford Nanopore Technologies) sequencing was carried out using Ligation Sequencing Kit (SQK-LSK110) and Native Barcoding Kit 96 (SQK-NBD112.96) according to manufacturers’ protocols. Before library preparation, DNA was quantified using the Quantus system (Promega Corp.). The integrity of samples was analyzed with TapeStation 2200 (Agilent). The library was sequenced using R9.4.1 MinION Flow Cell (FLO-MIN106D) for 72 hours ^77^. Base-calling and demultiplexing were carried out using the MinKnow software suite using the Super Accuracy option (www.nanoporeteflych.com). Contigs were assembled by a hybrid approach where long reads were initially used to generate a whole closed chromosome using the wf-bacterial-genomes workflow from EPI2ME Labs (https://github.com/epi2me-labs/wf-bacterial-genomes). This workflow uses flye software (https://github.com/fenderglass/Flye) for assembling and medaka software (https://github.com/nanoporetech/medaka) for long read-based contig polishing. In the next step, contigs were further polished with short reads using three tools: Polypolish (https://github.com/rrwick/Polypolish), Pilon (https://github.com/broadinstitute/pilon), and POLCA ^78^. Mutations detected in the genomes of the phage-resistant mutants were mapped against the *D. solani* WT reference genome ^29^. Genes found to have changes in their nucleotide sequence were characterized for their transcriptional organization using Operon-mapper (https://biocomputo.ibt.unam.mx/operon_mapper/) ^57^. Analyses of the biochemical pathways in which the selected mutated proteins might participate were done using KEGG ^79^. Similarly, mutated proteins were assessed for their possible roles in metabolic and functional cellular networks using STRING (Search Tool for Retrieval of Interacting Genes/Proteins) v11.5 (https://string-db.org/) (parameters: network type: *full network*, network edges: *high confidence*, interaction sources: *text mining, experiments, databases, co-expression, co-occurrence, gene fusion* ^42^), providing essential information concerning interactions of proteins of interest ^80^ using the proteome of *D. solani* strain IPO 2222 as a reference.

### Adsorption of ΦD5 to D. solani IPO 2222 wild-type and phage-resistant mutants

The rate of ΦD5 adsorption to *D. solani* IPO 2222 WT and phage-resistant mutants was determined as described before ^42^. Briefly, bacterial cultures in their log-phase growth were inoculated with a phage suspension (at a Multiplicity of Infection (MOI) of 0.01) and incubated for up to 20 min at 28 °C. Two individual samples of each bacterial strain tested were collected at various times: (0 (control), 1, 2, 5, 10, 15, and 20 minutes) ^42^, and the number of non-adsorbed phages was quantified. Bacteriophages suspended in sterile TSB medium and recovered at the same time points as above were used as a negative control. The experiment was repeated 3 times, and the results were averaged. Phage adsorption efficiency was calculated as described before ^42^.

### Assessment of morphological features of cells and colonies with microscopic techniques

As described previously, the colony morphology of phage-resistant mutants was analyzed with a Leica MZ10 F stereomicroscope with 10x and 40x magnifications coupled to a Leica DFC450C camera ^30, 57^. Likewise, the cell morphology of phage-resistant mutants was evaluated using transmission electron microscopy (TEM) ^31^. TEM analyses were done at the Laboratory of Electron Microscopy (Faculty of Biology, University of Gdansk, Poland). WT and phage-resistant mutants were adsorbed onto carbon-coated grids (GF Microsystems), directly stained with 1.5% uranyl acetate (Sigma-Aldrich), and visualized with an electron microscope (Tecnai Spirit BioTWIN, FEI) ^31^. At least ten images of each ΦD5-resistant bacterial variant and the WT strain were obtained to estimate cell diameter. For atomic force microscopic (AFM) analysis, bacteria were grown overnight in the M9 minimal medium supplemented with 0.4% glucose on a microscopy glass cover slide placed in a Petri dish, with gentle shaking (60 rpm) at 28 °C. The slides were washed with distilled water, and subsequently, samples were fixed with filtered 2.5% glutaraldehyde (Sigma Aldrich) for 2 hours, washed again, and air-dried. Cells were imaged using Bioscope Resolve (Bruker), in ScanAsyst (Peak Force Tapping) mode, with the application of ScanAsyst Air probe (f0 7.0 kHz, diameter <12 nm, k:0.4 N/m) ^81^. Post-imaging analysis and cell measurements (n= 12 to 20 cells) were performed using NanoScope Analysis 1.80 (Bruker).

### Isolation and visualization of lipopolysaccharide (LPS)

Lipopolysaccharides (LPS) of the WT and phage-resistant *D. solani* mutants were isolated with a Lipopolysaccharide Extraction Kit (Abcam, Symbios, Gdansk, Poland) as described previously ^57^. LPSs were separated utilizing 4–20% SDS-polyacrylamide gradient gel Mini-PROTEAN® TGX™ Precast Protein Gel, BioRad, Hercules, USA) electrophoresis (SDS PAGE) as described in ^82^ and visualized with silver staining as described before ^83^.

### Growth of the phage-resistant spontaneous mutants of D. solani wild-type strain in rich and minimal media

To determine whether phage resistance affects the growth rate of *D. solani* mutants, bacterial growth was assessed in both TSB (rich medium) and M9+0.4% glucose (minimal medium), as previously described ^31^. The experiment was replicated once, and the average generation time was determined using the Doubling Time Calculator (parameters: C0=3 h, Ct = 7 h, t= 4 h) (http://www.doubling-time.com/compute.php) ^84^. In addition, the ability of ΦD5-resistant mutants and the WT to grow at different temperatures was tested qualitatively on solid rich and minimal media incubated at 5, 8, 15, 28, and 37 °C, as described before ^85^. Growth was assessed visually on a daily basis. The experiment was repeated once using the same setup. To evaluate whether the ΦD5-resistance affects the growth rate of the mutants at different pHs, the growth rate of selected *D. solani* ΦD5-resistant mutants was compared in TSB at pH 4 and 10, similarly to other studies ^75^. In brief, overnight bacterial cultures in TSB (ca. 10^9^ CFU ml^-1^) were diluted 50-fold in fresh growth broth with pH 4 or 10. 100 µL of such prepared bacterial cultures were transferred to the wells of 96-well plates and wrapped with optically clear sealing tape (Sarstedt) to prevent drying up. The bacterial growth rate was determined by optical density (at 600 nm, OD_600_) measurements every 0.5 h for 16 h in an Epoch2 Microplate Spectrophotometer (BioTek). The experiment was repeated once, and the results were averaged. The generation time was calculated as described above.

### Evaluation of the phenotypes of phage-resistant bacterial mutants

The ability of phage-resistant *D. solani* mutants to use different carbon and nitrogen sources was analyzed in GEN III, EcoPlate, PM1, and PM2a 96-well plates in a BIOLOG phenotypic microarray system (Biolog Inc.) as described previously ^57^. In addition, the spontaneous ΦD5-resistant *D. solani* mutants were also screened for phenotypic features that may be crucial for their environmental fitness ^42^, including susceptibility to hydrogen peroxide ^75^, swimming and swarming motility ^75^, biofilm formation ^86^, the capability to grow on TSA medium containing 5% NaCl ^87^, production of enzymes: pectinolytic enzymes ^88^, cellulases ^89^, proteases ^90^ and siderophores ^91^. In addition, to test whether the phage resistance affects the profile of extracellular enzymes produced by *D. solani* IPO 2222, the phage-resistant mutants were tested, in duplicates, using API-ZYM stripes (bioMérieux) following the protocol provided by manufacturer ^30^.

### Ability of phage-resistant D. solani mutants to use sucrose, galacturonate, glucuronate, galactarate and pectin as a sole carbon source

Phage-resistant mutants were tested for the ability to metabolize different carbon sources released from plant tissues upon infection by *D. solani*. Overnight bacterial cultures grown in M9 medium + 0.4% glucose were diluted 50-fold in fresh M9 supplemented either with 0.4% sucrose, galacturonate, glucuronate, galactarate, or pectin (Sigma-Aldrich). One hundred microliters of diluted bacterial cultures were moved to sterile wells of the 96-well microtiter plates. The inoculated plates were closed with optically clear sealing tape (Sarstedt) to prevent contamination and evaporation of bacterial cultures. The growth rate was determined at 28 °C by measuring the optical density (OD600) every 0.5 h for a total time of 12 h in an Epoch2 Microplate Spectrophotometer (BioTek). Bacterial cultures in a 96-well plate were shaken (orbital shaker, 60 rpm) between the OD measurements to prevent anaerobic conditions and sedimentation of bacterial cells at the bottom of the well. The growth of each strain was analyzed in 6 technical replicates, and the results were averaged. Six wells inoculated with sterile growth medium served as a negative control, and the positive control was six wells inoculated with bacterial cultures grown in M9 medium + 0.4% glucose. The experiment was repeated once.

### Proteomics of phage-resistant D. solani mutants grown in rich and minimal media

#### Sample preparation

Mass spectrometry (MS) analysis of proteomes of *D. solani* wild-type strain and phage-resistant mutants was performed to verify whether the phage resistance affects the profile of total proteins produced by the resistant mutants compared to the WT strain. For that, bacteria were grown either in TSB (rich medium) or in M9 minimal medium supplemented with glucose (0.4% final concentration) (minimal medium) for 24 h at 28 °C with shaking (120 rpm). After this time, bacterial cells from a 20 ml culture were collected by centrifugation (8000 x g, 5 min.) and washed two times with PBS pH 7.2 buffer (Sigma-Aldrich). Washed bacterial cells were dissolved in lysis buffer (1% SDS, 100 mM Tris/HCl pH 8, 50 mM DTT) and incubated at 95 °C for 10 min to lyse the cells. After incubation, the lysates were cooled to room temperature and sonicated for 5 min (15-sec pulse, 10 s rest, 100% amplitude) using a QSonica Q500 sonicator (Cole-Parmer, Vernon Hills, IL, USA). Protein concentration was determined by measuring absorbance at 280 nm using μDrop plate of Multiskan GO (ThermoScientific, USA). For each sample, 100 µg of proteins were mixed with 200 µL of buffer containing urea (8 M urea in 100 mM Tris/HCl pH 8.5). Proteins were digested with MS Grade trypsin (Promega, Madison, WI, USA) on 10 kDa Microcons (Merck-Milipore, Burlington, MA, USA) according to a Filter-Aided Sample Preparation (FASP) procedure ^92^. Final clean-up was done on a C18 resin using the STAGE Tips procedure ^93^.

#### LS-MS/MS and SWATH analyses

The LC-MS/MS analyses were performed on an Ekspert MicroLC 200 Plus system (Eksigent, Redwood City, CA, USA) coupled with a hybrid TripleTOF 5600+ mass spectrometer with DuoSpray Ion Source (AB SCIEX, Framingham, MA, USA). Chromatographic separation was carried out on a ChromXP C18CL column (3μm, 120 Å, 150×0.3 mm; Eksigent). The chromatographic gradient for the data-dependent acquisition (DDA) and SWATH analyses was 11-42.5% B (solvent A: 0% aqueous solution. 0.1% formic acid; solvent B: 100% acetonitrile, 0.1% formic acid) with a flowrate of 10μL/min for 60 min. All experiments were performed in a positive ion mode. The system was controlled by SCIEX Analyst TF 1.7.1 software. Each cycle of the applied DDA method comprised precursor spectra accumulation in 100 ms in the range of 400-1200 m/z, followed by top 20 candidate ions per scan in 50 ms in the range of 100-1800 m/z resulting in a total cycle time of 1.15s. Experiments of triplicated samples were performed in a looped product ion and high sensitivity modes. A set of 25 transmission windows of variable width was constructed with the SWATH variable window calculator, covering the precursors’ mass range of 400-1200 m/z. The collision energy for each window was calculated for +2 and +5 charged ions centered upon the window with a spread of five. The SWATH-MS/MS survey scan was acquired in the range of 400-1200 m/z with an accumulation time of 50 ms, followed by SWATH-MS/MS spectra scans in the range of 100-1800 m/z during 40 ms accumulation time resulting in the total cycle time of 1.1 s.

#### Database search and data analysis

Database search was conducted with ProteinPilot 5.0.2 software (AB SCIEX) using the Paragon algorithm against the reviewed *Dickeya solani* database (18.11.2022, Uniprot (https://www.uniprot.org/)) with the following parameters: instrument TripleTOF5600, alkylation of cysteines by iodoacetamide, trypsin enzyme digestion, ID focus on biological modifications, search effort ‘through ID’, and detected protein threshold [Conf] > 10%, with an automatic false discovery rate (FDR) analysis. Spectral library was created in PeakView 2.2 software (AB SCIEX) using set parameters: a maximum of 6 peptides per protein and 6 transitions per peptide; peptides with the confidence of at least 95% and an extraction window width of 15 min and 75 ppm XIC width. All results were exported to MarkerView software (AB SCIEX) and normalized using the total area sums (TAS) approach. Statistical analysis of the outcome was performed in Perseus 2.0.7.0 software ^94^, where second normalization (log(2x)) was carried out. A two-sample *t*-test was performed to obtain q-values for each listed protein (adjusted p-value; q-value <0.05). In addition, fold-change (FC) values were calculated to collect information about the up- and down-regulation of each protein. Outcome visualization was generated with the InteractiVenn tool (http://www.interactivenn.net/). The mass spectrometry proteomics data have been deposited to the ProteomeXchange Consortium (http://proteomecentral.proteomexchange.org) via the PRIDE partner repository ^95, 96^ with the dataset identifier PXD038825.

#### Sensitivity of the phage-resistant D. solani mutants to surfactants

To determine whether the phage resistance affects the bacterial cell surface stability, the growth of the selected *D. solani* ΦD5-resistant mutants was assessed at 28 °C for 16 h in TSB supplemented with selected surfactants (0.1% Triton X-100, 0.1% Tween 20, 0.1% Tween 80, 0.1% Poloxamer 407, 0.1% N-lauroyl sarcosine or 0.1% EDTA) (Sigma-Aldrich) as previously described ^31^. The average generation time of each mutant was analyzed using the Doubling Time calculator (parameters: C0= 3 h, Ct=7 h, t=4 h) (http://www.doubling-time.com/compute.php) ^84^ as described above. *D. solani* wild-type strain IPO 2222 was used as a control. The experiment was repeated once, and the results were averaged.

#### Autoaggregation of phage-resistant D. solani mutants

The autoaggregation of phage-resistant *D. solani* mutants was measured as enhanced sedimentation and phase separation of aqueous bacterial suspensions, as previously described ^97^. Phage-resistant mutants were evaluated for the rate of autoaggregation (sedimentation) as described in ^56, 98^. Briefly, the WT strain and ΦD5-resistant mutants were grown in 10 ml of TSB at 28 °C with shaking (120 rpm) for 24 h, and 1 ml of bacterial culture was then transferred to sterile cuvettes (Eppendorf). The optical density (OD_600_) was initially measured (time = 0 h), and after incubation at 28 °C for 24 h. Each phage-resistant bacterial mutant was analyzed in duplicates, the experiment was repeated once, and the results were averaged ^57^. Ratio aggregation (sedimentation) was assessed as follows: %A = 1-(OD_600_ 24 h/OD_600_ 0 h), where: %A—the percentage of aggregation (sedimentation), OD_600_ 0h—OD of bacterial culture at time 0 h, OD_600_ 24 h— OD of bacterial culture at time 24 h ^98^.

#### Antibiotic susceptibility of the phage-resistant mutants of D. solani IPO 2222

The antibiotic susceptibility of phage-resistant mutants was determined as previously described ^57, 99^. Disks containing antibiotics (BD BBL - Sensi-Disc antimicrobial test discs): ciprofloxacin (5 µg), chloramphenicol (30 µg), ceftazidime (30 µg), linezolid (10 µg), gentamicin (10 µg), aztreonam (30 µg), tigecycline (15 µg), doxycycline (30 µg), imipenem (10 µg), vancomycin (5 µg), erythromycin (15 µg), streptomycin (300 µg) were placed on Mueller-Hinton (MH medium, BD) supplemented with 1.5 % agar (Oxoid) inoculated with bacteria. Before testing antibiotic susceptibility, phage-resistant mutants and the WT strain were grown for 16 h in TSB with shaking (120 rpm). Plates inoculated with bacteria and antibiotic disks were cultured at 28 °C for 24 h and subsequently observed for a clear halo (indicating growth inhibition zone) surrounding a given disc indicating antibiotic susceptibility. The experiment was repeated once.

#### Virulence of phage-resistant D. solani IPO 2222 mutants on chicory leaves

The ability of ΦD5-resistant bacterial mutants to cause maceration of chicory leaves was assessed as previously described ^100^. Briefly, chicory leaves (5 replicate leaves per bacterial strain) were purchased locally in Gdansk, Poland. The leaves were inoculated with a given bacterial strain by creating a shallow, ca. 1-cm-long cut across each leaf with a sterile scalpel and subsequently inoculating with 10 µL of bacterial suspension (ca. 10^8^ colony forming units (CFU) mL^-1^). Inoculated chicory leaves were sealed in a plastic bag containing filter paper moistened with sterile water. After incubation for 48 h at 28 °C, the diameter of the rotted tissue on each inoculated leaf was measured ^100^. WT *D. solani* was used as a positive control, and negative control tubers were inoculated with sterile demineralized water. The experiment was repeated once, and the results were averaged.

#### Virulence of phage-resistant D. solani mutants on potato plants grown in potting soil

Experiments involving plants grown in a growth chamber were performed using the previously developed protocol ^71^. Certified potato tubers (cv. Kondor) were purchased from the Plant Breeding and Acclimatization Institute (National Research Institute, Poland) and were cultivated as described earlier^71^. After two weeks, rooted plants (ca. 10–15 cm in height) were transferred to 1 L pots and grown in potting soil for additional two weeks. Potato plants (5 plants per repetition) were inoculated with phage-resistant *D. solani* mutants or the WT strain by the application of 50 ml of bacterial suspensions (ca. 10^8^ CFU mL^−1^) in sterile Ringer’s buffer directly to the soil surrounding the base of stems 1 h after plants had been well irrigated to guarantee uniformly moist soil. As a negative control, the soil was treated with sterile Ringer’s buffer (50 ml per plant) alone. Pots were then randomized in a growth chamber with 4 blocks of 10 pots. Plants were visually examined daily for the development of disease symptoms (chlorosis, black rotting of the stem, haulm wilting, plant death). Plants were sampled for bacterial populations 14 days after inoculation. Per the analyzed plant, stem sections (ca. 2 cm long) located ca. 5 cm above ground level were collected, combined into one sample, and surface-sterilized as described before ^57^. The number of bacterial cells within potato stems was determined by dilution-plating stem macerates on CVP medium supplemented with 200 μg mL^−1^ cycloheximide and counting the resulting cavity-forming *D. solani* colonies. The experiment was repeated once, and the results were averaged for analysis.

#### Statistical analysis

Statistical analyses were carried out using a previously described approach ^42^. To achieve normality, bacterial colony counts were transformed to log(x + 1) ^101^. Treatments were examined following the experimental design, which, each time, involved doing two independent duplicated experiments for every treatment applied. For samples in which bacterial population sizes were normally distributed, the Shapiro-Wilk test (at p=0.05) was applied ^102^. The Welch’s T-test was used for samples in which population sizes were not normally distributed, such as: (i) control vs. treatment or (ii) treatment vs. treatment ^103^. The Fisher-Snedecor test was used to validate the homogeneity of variance ^104^. A two-tailed Student’s t-test was used to assess pairwise differences between analyzed samples ^105^. The linear model included an entire block design, and replicates were treated as separate blocks ^106^. The impact of time, the treatment, and the two-way interaction between the time and treatment type were examined in the model.

## Acknowledgments

This research was financially supported by the National Science Center, Poland (Narodowe Centrum Nauki, Polska) via a research grant OPUS 13 (2017/25/B/NZ9/00036) to Robert Czajkowski. The authors would like to express their gratitude to Prof. Steven E. Lindow (University of California-Berkeley, Berkeley, CA, United States) for his comments on the manuscript and his editorial work and Prof. Stanislaw Oldziej (University of Gdansk, Gdansk, Poland) for helpful discussions about protein structures and interactions.

## Author contributions

Conceptualization: RC, Methodology: DS, LR, MK, MN, IM, PC, SJ, Investigation: DS, AS, PB, LR, MK, MN, DMK, MR, IM, SJ, Writing-original draft preparation: RC, LR, IM, PC, Writing-Review & Editing: RC, SJ, PC, Visualization: MN, PC, IM, MR, DMK, SJ, AS, Funding acquisition: RC. All authors reviewed the manuscript before submission.

## Data availability

Data generated or analyzed during this study are included in this published article and its supplementary materials. Correspondence and requests for materials should be addressed to R.C. In addition, the mass spectrometry proteomics data have been deposited to the ProteomeXchange Consortium *via* the PRIDE partner repository with the dataset identifier PXD038825. The DsR34 and DsR207 complete genomes were deposited in the NCBI GenBank database under accession numbers CP110886 and CP110887, respectively. Raw TEM and AFM photos and accompanying data are deposited at Zenodo (www.zenodo.org) under https://doi.org/10.5281/zenodo.7550485.

