## Supplementary Dataset 1 for "Being spontaneous has its costs! Characterization of spontaneous phage □D5-resistant mutants of *Dickeya solani* strain IPO 2222"

**Supplementary Figure 1.** Function category of significantly differentially abundant proteins in phage-resistant mutants DsR34 and DsR207 and the wild-type IPO 2222 in TSB (rich medium) (**A**) and M9+glucose (minimal medium) (**B**)

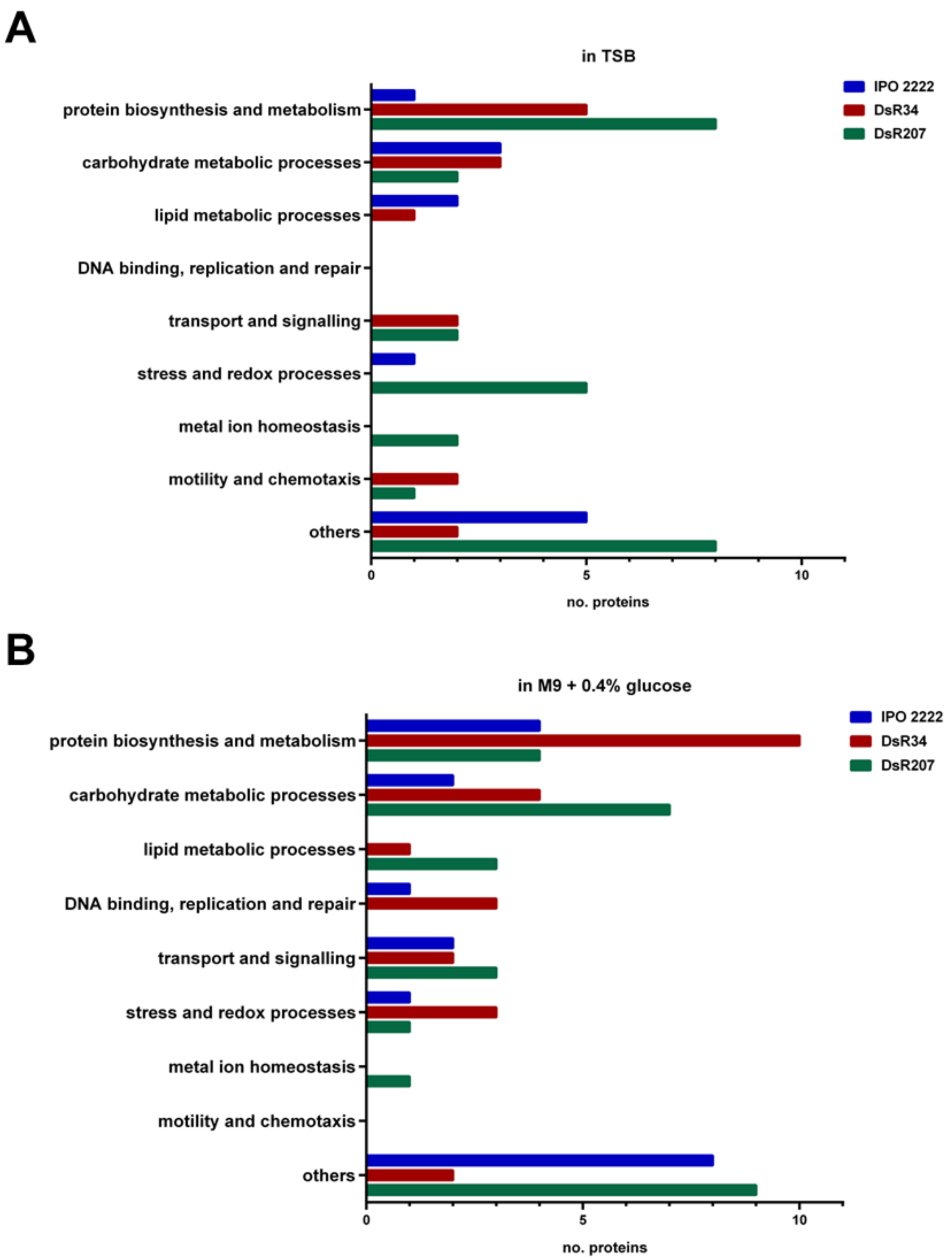
